# Spatially resolved translational dysregulation in *Grin2a*+/- mouse model of schizophrenia

**DOI:** 10.64898/2025.12.23.696217

**Authors:** Mingrui Wu, Jiahao Huang, Sameer Aryal, Zohreh Farsi, Hongyu Chen, Yiming Zhou, Shuchen Luo, Wendy Xueyi Wang, Kevin Bonanno, Margaret Yin, Inès Picard, Borislav Dejanovic, Hasmik Keshishian, Steven A. Carr, Morgan Sheng, Xiao Wang

## Abstract

Loss-of-function (LoF) mutations of *GRIN2A*, encoding the GluN2A subunit of N-methyl-D-aspartate receptor (NMDAR), confer a high risk for schizophrenia (SCZ)^1–3^, yet how they affect diverse brain cell types remains poorly understood. Here, we combined subcellular-resolution spatial omics technologies, STARmap^4^ and RIBOmap^5^, to jointly resolve single-cell transcriptomes and translatomes for 3,447 genes in the brains of *Grin2a*+/- mice and their wild-type littermates across 538,188 cells. Translational dysregulation was markedly more prominent than transcriptional changes in neurons. Across neuronal subtypes, a set of genes including *Camk2a*, *Arc*, *Egr1*, *Egr3*, *Chmp2b*, and *Pja2* exhibited translational reduction in a *Grin2a* gene dose-dependent fashion, suggesting a connection between NMDAR hypofunction and reduced protein synthesis of downstream synaptic plasticity effectors. In interneurons (particularly parvalbumin interneurons), a strong reduction of *Gad2* translation implies loss of inhibitory function in cortical microcircuits, which has long been hypothesized for SCZ pathophysiology. Non-neuronal cell types including astrocytes, oligodendrocytes, and vascular cells also exhibited region-specific translational changes in neurotransmitter transport, lipid synthesis, myelination, and stress response pathways, some of which co-varied with regional neuron state. Together, our study reveals brain-wide translation dysregulation as a critical mechanism underlying SCZ pathophysiology.

## Introduction

Schizophrenia (SCZ) is a chronic and severe neuropsychiatric disorder affecting approximately 1% of the global population^6,7^. It is characterized by a constellation of symptoms, including delusions, hallucinations, cognitive impairments, and social withdrawal^1,8^. Despite decades of research, the molecular and neurobiological basis for SCZ remains poorly understood, while current pharmacological treatments, primarily targeting the dopaminergic signaling pathway, offer only partial relief to symptoms and are ineffective for many patients^1^.

SCZ has a strong genetic basis (heritability 60-80%)^9–11^, with hundreds of associated common variants of small effect identified through genome-wide association study (GWAS)^2,12^. Recent exome sequencing study from the Schizophrenia Exome Sequencing Meta-analysis (SCHEMA) consortium further revealed risk genes with rare loss-of-function (LoF) variants that confer substantial risk for SCZ^3,13^. Many of these SCZ risk genes converge on pathways related to synapse formation, structure, and function, with particular enrichment in glutamatergic neurotransmission^13^.

*GRIN2A*, encoding the GluN2A subunit of the N-methyl-D-aspartate receptor (NMDAR)^14^, has been repeatedly identified as a risk gene for SCZ in large-scale common^2^ and rare variant^3^ human genetics studies. This association between *GRIN2A* LoF and SCZ is in line with the hypoglutamatergic hypothesis of SCZ, which posits that dysfunction of glutamatergic neurotransmission contributes to SCZ pathophysiology^15–17^. Notably, *Grin2a* heterozygous mutant mice show EEG abnormalities and other neurobiological features that resemble those observed in humans with SCZ^18,19^. Transcriptomic characterization of *Grin2a*+/- mice revealed widespread alterations in activity-, mRNA translation-, and metabolism-related genes across different brain regions^19^. However, transcriptome profiling alone cannot inform whether observed changes in mRNA reflect alterations in protein abundance—a critical gap given that mRNA levels correlate poorly with protein expression and that translational dysregulation has emerged as a key mechanism in neurodevelopmental and psychiatric disorders^20–24^. Direct measurement of actively translated mRNAs is therefore essential to determine the extent to which *Grin2a*+/-contributes to SCZ pathophysiology via translational alterations.

To address this question, we employed both STARmap and RIBOmap to create an integrative spatial map of molecular changes in the *Grin2a*+/- mouse brain with micron-level spatial precision. STARmap enables spatially resolved quantitation of mRNA transcripts *in situ*, while RIBOmap extends this capability by selectively measuring ribosome-associated mRNAs, thereby dissecting transcriptional versus translational alterations. By integrating these data layers, we clarify the cell-type- and region-specific effects of *Grin2a* LoF, providing unprecedented insights into the cellular and circuit pathophysiology underlying the *Grin2a*+/- model of SCZ.

## Results

### Spatial transcriptional and translational co-profiling of *Grin2a*+/- mouse brain

We applied STARmap and RIBOmap to profile 3,447 genes of interest in brains from *Grin2a*+/-mice and their wild-type littermates at 12 weeks of age (**Fig. 1a, Supplementary Table 1**). The 3,447-gene panel was compiled from a pool of canonical brain cell-type markers, along with genes shown to be differentially expressed in bulk RNA-seq, single-nucleus RNA-seq (snRNA-seq), or proteomic profiling of *Grin2a* mutant mice^19,25^. We sampled the brain at three coronal positions along the anterior-posterior axis and, at each position, collected two adjacent 20-μm slices to be processed with STARmap and RIBOmap, respectively (**Fig. 1a**). In total, we profiled 24 brain slices, which together included 538,188 cells (**Extended Data Fig. 1a,b**) and covered multiple brain regions implicated in SCZ^26–28^, such as prefrontal cortex (PFC), somatosensory cortex (SSC), striatum (ST), hippocampus (HP), and thalamus (TH).

**Fig. 1:**
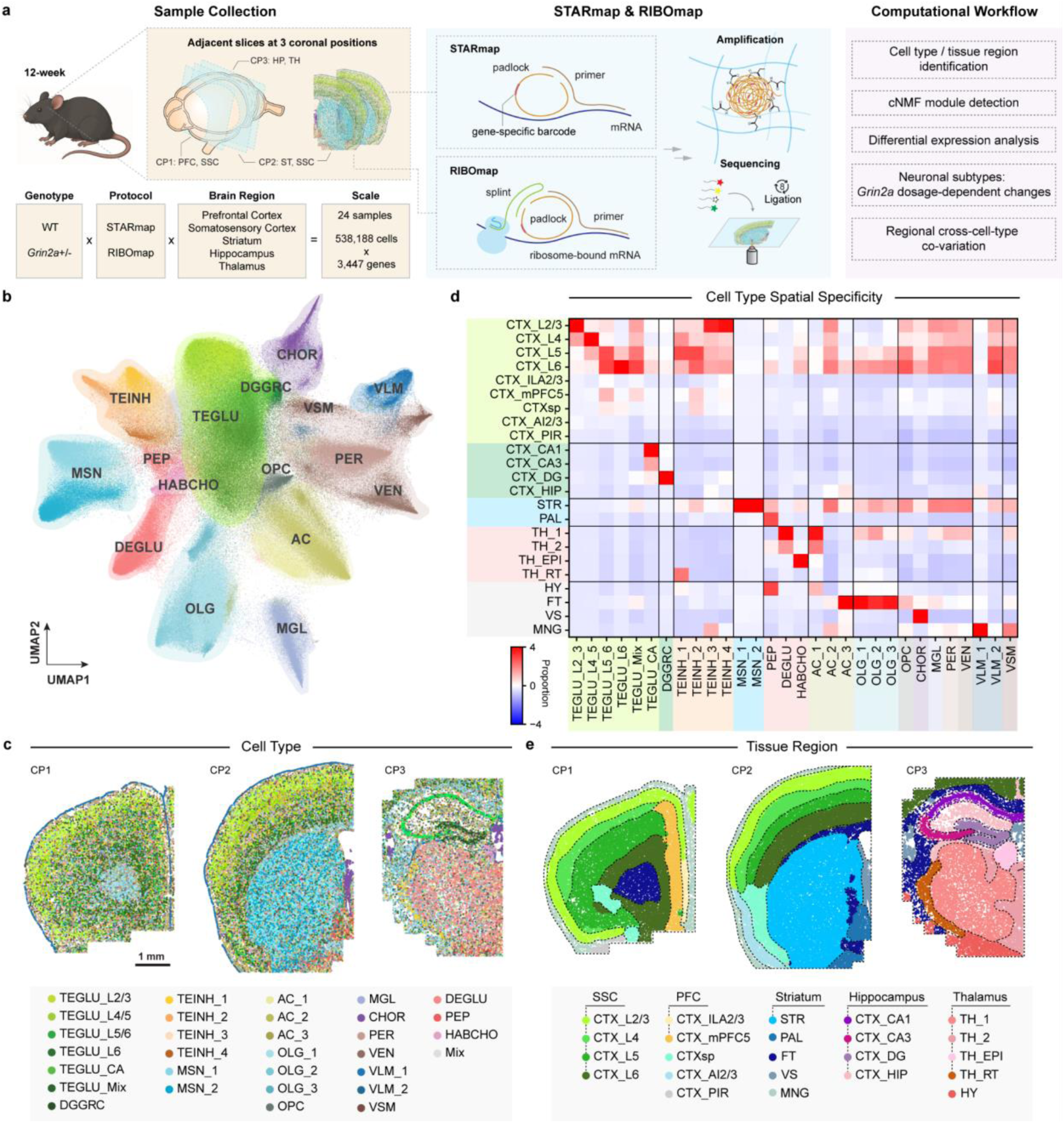
Integrative profiling of *Grin2a*+/- mouse brain. **a**, Research schematics. Mouse brain slices were collected from 12-week wild-type and *Grin2a*+/-mice at three coronal positions, covering multiple brain regions of interest: prefrontal cortex (PFC), somatosensory cortex (SSC), striatum (ST), hippocampus (HP), and thalamus (TH). RIBOmap and STARmap targeting 3,447 genes were performed on each pair of adjacent slices, respectively. Computational workflow included cell type classification, tissue region identification, differential expression (DE) analysis, and cross-cell-type co-variation analysis. **b**, Uniform Manifold Approximation and Projection (UMAP) of 538,188 cells colored by detailed subtypes and labeled with top-level major cell types: AC, astrocytes; CHOR, choroid plexus epithelial cells; DEGLU, diencephalon excitatory neurons; DGGRC, dentate gyrus granule cells; HABCHO, habenula cholinergic neurons; MGL, microglia; MSN, medium spiny neurons; OLG, oligodendrocytes; OPC, oligodendrocyte precursor cells; PEP, peptidergic neurons; PER, pericytes; TEGLU, telencephalon projecting excitatory neurons; TEINH, telencephalon inhibitory interneurons; VEN, vascular endothelial cells; VLM, vascular and leptomeningeal cells; VSM, vascular smooth muscle cells. **c**, Spatial visualizations showing subtypes distribution at 3 coronal positions. **d**, Heatmap showing cell subtype compositions across molecular tissue regions. The proportion of each cell subtype is calculated for every molecular tissue region. Then, the z-scores of these percentages are plotted for each cell type, with subtypes grouped under their respective top-level cell types. CTX, Cerebral cortex; ILA, Infralimbic area; CTXsp, Cortical subplate; AI, Agranular insular area; PIR, Piriform area; HIP, Hippocampal region; CA, Ammon’s horn; DG, Dentate gyrus; STR, Striatum; PAL, Pallidum; TH, Thalamus; EPI, Epithalamus; RT, Reticular nucleus; HY, Hypothalamus; FT, Fibre tracts; VS, Ventricular systems; MNG, Meninges. **e**, Spatial visualizations showing the delineation of molecular tissue regions by SPIN at 3 coronal positions.

To generate cell-type annotations for downstream analysis, we first integrated the single-cell transcriptomic and translatomic profiles with 1,017 canonical cell type markers ( **Extended Data Fig. 1c,d**) and identified 16 major cell types and 30 subpopulations through unsupervised Leiden clustering (**Fig. 1b,c**). Clusters were annotated according to the spatial mouse CNS atlas by Shi *et al*^25^. In detail, we resolved 7 major neuronal cell types, including telencephalon excitatory neuron (TEGLU), diencephalon excitatory neuron (DEGLU), telencephalon inhibitory neuron (TEINH), and medium spiny neuron (MSN), and 9 major non-neuronal cell types, including astrocyte (AC), oligodendrocyte (OLG), microglia (MGL), and pericyte (PER) (**Extended Data Fig. 2a, 3a**). Their respective subclusters correspond to distinct spatial localization patterns (**Fig. 1c**) and functional roles. For example, TEGLU subtypes separated spatially into cortical layers and the Cornu Ammonis (CA) in the hippocampus, while two MSN subclusters, both localized in the striatum, respectively represent direct-pathway and indirect-pathway spiny projection neurons (dSPNs and iSPNs) (**Extended Data Fig. 2b**).

To assess spatial heterogeneity across and within brain regions, we identified 23 molecular tissue regions using SPIN^29^ (**Fig. 1d,e, Extended Data Fig. 3b,c**), including cortical layers (CTX_L2/3, CTX_L4, CTX_L5, CTX_L6), PFC regions (CTX_ILA2/3, CTX_mPFC5), hippocampal fields (CTX_CA1, CTX_CA3, CTX_DG, CTX_HIP), and thalamic compartments (TH_1, TH_2, TH_EPI, TH_RT). Notably, the resolution of our method was sufficient to delineate fine tissue structures. For instance, the thalamic reticular nucleus (TRN) emerged as a discrete domain from TH_1 and TH_2 (**Fig. 1e**). The TRN (TH_RT) is challenging to isolate by anatomical dissection, yet is crucial for regulating thalamocortical signaling and a key node implicated in SCZ pathophysiology^30,31^. Cell-type enrichment across molecular tissue regions was concordant with current neurobiological understandings. For example, MSN and DEGLU were predominantly enriched in the striatum (STR) and the thalamus (TH_1, TH_2), respectively (**Fig. 1c, Extended Data Fig. 3d**). Together, these labels provide a basis for identifying disease-associated changes that localize to specific cell types and brain regions.

### RIBOmap reveals global translational reduction of synapse- and translation-associated gene modules

To gain an overview of transcriptional and translational changes in *Grin2a*+/- animals, we applied consensus non-negative matrix factorization (cNMF)^32^ to identify gene modules with distinct expression patterns across the entire dataset (**Fig. 2a, Extended Data Fig. 4a,b**). Among the 35 modules identified, 24 showed cell-type-specific enrichment (M1-M24), while 11 were not confined to particular cell types (M25-M35) (**Extended Data Fig. 4c,d, Supplementary Table 2**). Differential expression analysis of cNMF modules revealed greater downregulation in RIBOmap compared to STARmap, particularly for M2, M4, M7, M27, M28, M29, and M33, indicating potential reduction at the translational level for these modules in *Grin2a*+/- animals (**Fig. 2a, bottom two rows**). We also noticed a higher number of differentially expressed genes (DEGs) detected by RIBOmap than by STARmap for most major cell types, further highlighting dysregulation of translational states not obvious at the mRNA level (**Fig. 2a right**). To identify the biological processes associated with each cNMF module, we performed Gene Ontology (GO) enrichment analysis on the top 50 genes within each module^33^ (**Fig. 2b**). As anticipated, cell-type-specific modules were enriched for canonical functions of their respective cell types, such as glutamatergic synaptic transmission (M1) in TEGLU, neuropeptide activity (M5) in TEINH, microglia cell activation and synapse pruning (M17) in MGL; while non-cell-type-specific modules showed broader functional enrichments such as postsynaptic density (M28) and translation (M33).

**Fig. 2:**
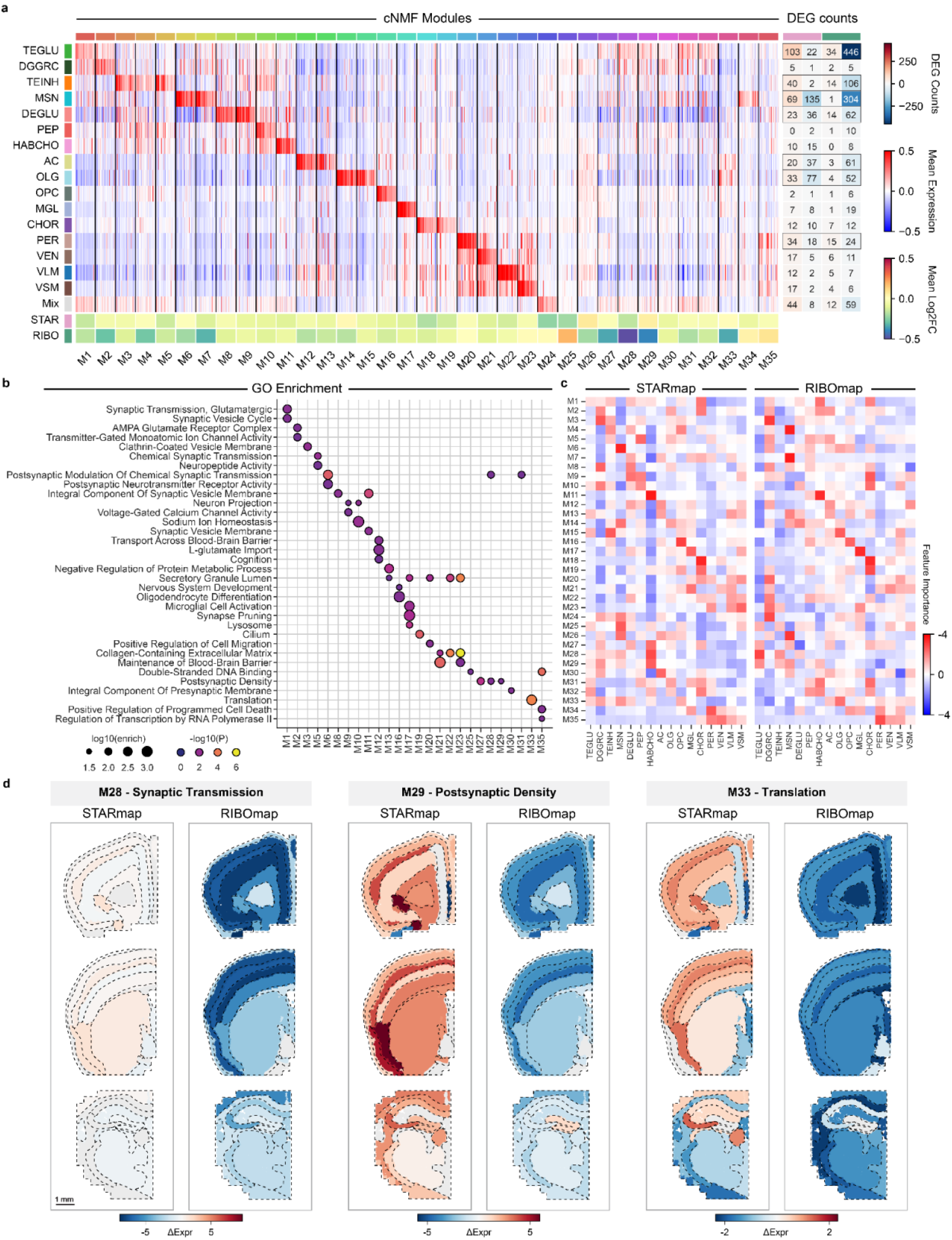
System-level analysis of gene expression modules in transcriptome and translatome. **a**, Gene expression module identification with cNMF in spatial transcriptome and translatome. Multi-panel heatmap showing 35 detected gene modules: Expression of top 50 genes with the highest loading of each module in major cell types (main), cell-type-specific DEG count (right), and averaged log fold-change in STARmap and RIBOmap of top-50 genes from each module in the total cell population (bottom). Cell types showing prominent changes were outlined in black. **b**, GO enrichment analysis of cNMF modules with the top 50 genes; Color scale, negative log-transformed adjusted p-value; dot size, log-transformed enrichment score. **c**, Heatmap displaying module importance for predicting genotypes across major cell types in STARmap (left) and RIBOmap (right). A Random Forest classifier was trained using aggregated module expression to distinguish WT from *Grin2a*+/− cells. **d**, Spatial visualizations of differential expression of module M28, M29, and M33 in STARmap (left) and RIBOmap (right).

To identify cNMF modules that explain the most difference between WT and *Grin2a*+/- mutant, we trained a Random Forest classifier to predict genotype labels within major cell types and ranked modules based on their feature importance (**Fig. 2c**). Notably, M28 and M29 were key genotype-associated features of TEGLU, TEINH, MSN, and DEGLU populations in RIBOmap, while M31-33 effectively classified TEGLU in STARmap. Additionally, M35 emerged as a top genotype-associated feature for vascular populations (PER, VEN, VLM) across both datasets. We next aimed to gain neurobiological insights from region-specific changes of these genotype-associated modules. Two broad observations emerged (**Fig. 2d**): First, module-level gene expression changes demonstrated spatial heterogeneity not only among major brain structures but also within subregions, highlighting the detailed regulatory signatures that bulk methods overlook. Second, modules displayed distinct patterns at the transcriptomic and translatomic levels. In detail, synaptic function-related modules (M28, M29) were generally downregulated at the translational level (RIBOmap) across most regions, but upregulated at the mRNA level (STARmap) in areas including CTX_L4, CTXsp, CTX_ILA2/3, and STR. Additionally, translation-related module M33 showed downregulation in prefrontal cortex (CTX_mPFC5) and slight upregulation in other cortical layers, striatum, and hippocampus in STARmap, consistent with previous snRNA-seq findings in *Grin2a* LoF mutants^19^. In contrast, M33 was downregulated across the entire cortex in RIBOmap, particularly in CTX_L5, CTX_mPFC5, and fiber tracts (FT). Such a discrepancy between STARmap and RIBOmap is indicative of translational regulation, as supported by global downregulation of the translation-related module M33, which might result in a net reduction in protein synthesis despite rising mRNA levels. This integrative analysis of gene modules showed that neuronal populations and many gene modules, particularly those associated with synaptic functions, exhibited greater translational than transcriptional alterations.

### *Grin2a* LoF triggers distinct molecular responses in major neuronal populations

To dissect single-gene alterations upon *Grin2a* LoF in neurons, we first examined the primary excitatory and inhibitory neuronal cell types in our dataset: TEGLU, DEGLU, TEINH, and MSN (**Fig. 3a-e**). These four cell types exhibited relatively high *Grin2a* expression (**Extended Data Fig. 5a**) and showed extensive molecular changes at both transcriptional and translational levels.

**Fig. 3:**
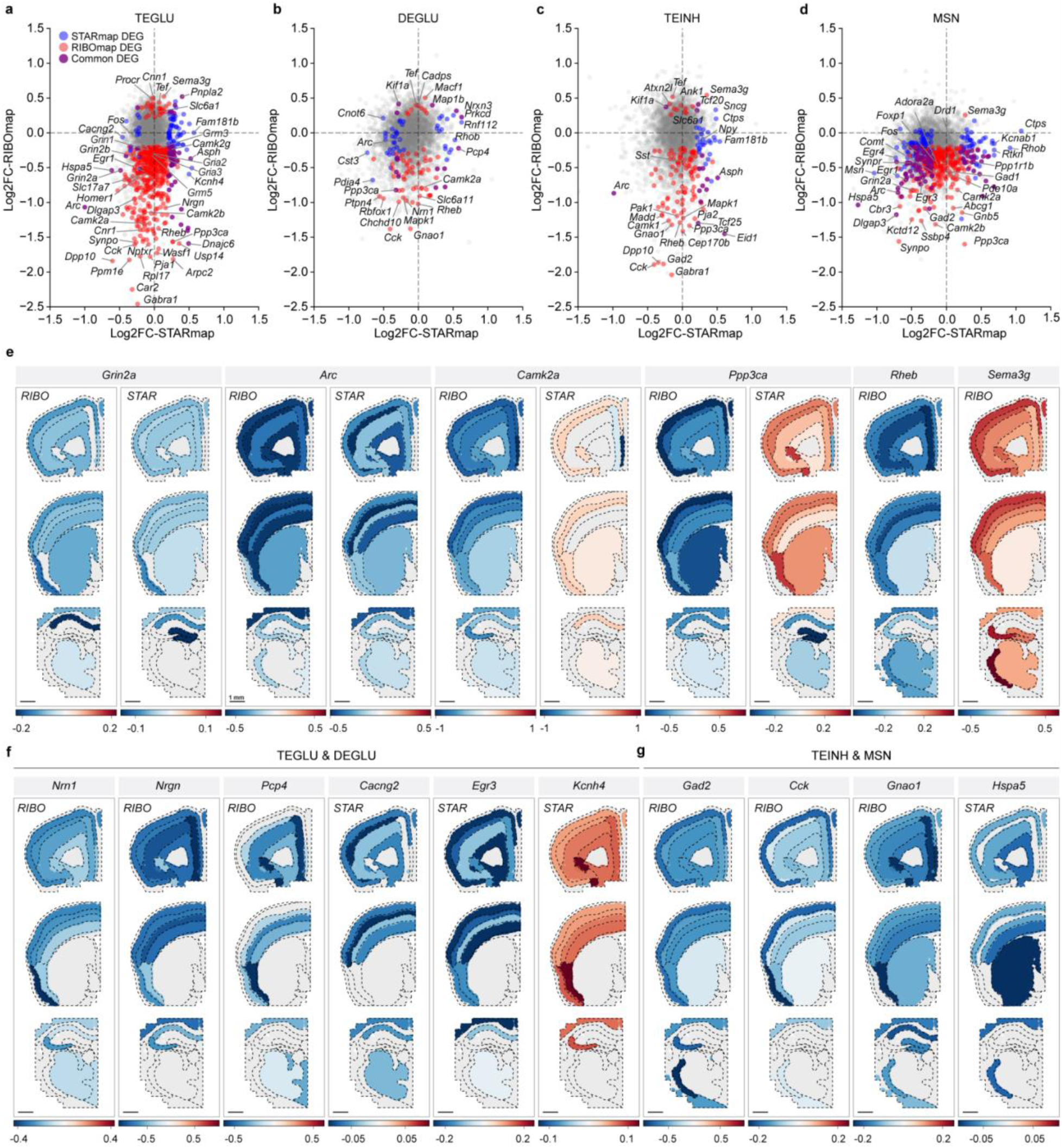
*Grin2a* LoF triggers distinct molecular responses in major neuronal populations. **a-d**, STARmap vs. RIBOmap log fold-change plots of 4 neuronal cell types of interest: TEGLU (**a**), DEGLU (**b**), TEINH (**c**), and MSN (**d**). DEGs with significant (adjusted p-value < 0.05, |Log2FC| ≥ 0.2) log fold-change in STARmap only, RIBOmap only, and both modalities are colored in blue, red, and purple, respectively. **e,** Spatial visualizations of representative differentially expressed genes in four neuronal populations of interest. **f, g**, Spatial visualizations of differentially expressed genes for excitatory neurons TEGLU and DEGLU (**f**) and inhibitory neurons TEINH and MSN (**g**) identified in STARmap and RIBOmap. Regional log-fold changes were calculated for the neuronal populations shown in each panel.

The prominent impact on TEGLU and DEGLU is consistent with the primary role of NMDAR in excitatory glutamatergic neurotransmission: GluN2A-containing NMDARs are predominantly localized to excitatory synapse, where they mediate calcium-dependent signaling and synaptic plasticity essential for learning and memory^34,35^. Despite shared disruption of synaptic plasticity genes (*Camk2a*, *Ppp3ca, Arc*), TEGLU and DEGLU exhibited markedly different profiles of glutamatergic pathway alterations, with TEGLU showing substantially more extensive dysregulation (**Fig. 3a,b,f**). In cortical TEGLU, the impact of *Grin2a* haploinsufficiency was much stronger in RIBOmap (translatome) than in STARmap (transcriptome). *Grin2a*+/- was associated with concurrent RIBOmap downregulation of *Grin1*, encoding the obligatory NMDA receptor subunit, and *Grin2b*, encoding the alternative GluN2B subunit, suggesting a general lessening of NMDA receptor-mediated signaling rather than compensatory subunit switching. Moreover, translational downregulation extended to AMPA receptor subunits (*Gria2*, *Gria3*), metabotropic glutamate receptors (*Grm5*, *Grm3*), and vesicular glutamate transporter *Slc17a7* (**Fig. 3a,f, Supplementary Table 3**). We also found widespread downregulation of glutamatergic postsynaptic scaffolds (*Dlg1*, *Dlg2*, *Dlgap3*, *Shank2*) and synaptic adhesion molecules (*Nrxn1*, *Nrxn2*) by RIBOmap. Downstream signaling components (*Camk2b*, *Camk2g*, *Homer1*) and some activity-regulated genes (*Egr1*, *Egr3*, *Nrn1*, *Arc, Nptxr*) were also reduced in RIBOmap (**Fig. 3f, Supplementary Table 3**). These changes collectively indicate failure of glutamatergic neurotransmission and attenuated synaptic plasticity in cortical excitatory neurons, which may contribute to SCZ-associated cognitive deficits. Compared with cortical TEGLU neurons, thalamic DEGLU exhibited fewer translatomic changes, with downregulation affecting genes involved in calcium-dependent signaling (*Camk2a*, *Ppp3ca*, *Pcp4*), while activity-regulated genes were relatively unperturbed (**Fig. 3b,f**).

Excitatory-inhibitory (E/I) balance is critical for proper cortical circuit function and cognition, and its disturbance has been repeatedly implicated in SCZ pathophysiology^36–38^. Studies of postmortem brain tissue from SCZ patients have revealed reduced expression of various GABAergic markers, including GAD1 (also known as GAD67) and parvalbumin (PV), particularly in cortical interneurons^39,40^. Such findings have led to the idea that dysfunction of cortical microcircuits involving both inhibitory interneurons and excitatory pyramidal neurons underlies the cognitive deficits of SCZ. In our spatial RIBOmap analysis, all four TEINH subtypes: TEINH_1-[*Pvalb*_*Gad1*], TEINH_2-[*Sst*_*Npy*], TEINH_3-[*Lamp5*_*Npy*], and TEINH_4-[*Vip*_*Cnr1*], showed marked reduction in translation of *Gad2*, with the largest magnitude of effect (|Log2FC| > 2) in PV interneurons (TEINH_1-[*Pvalb*_*Gad1*]) (**Fig. 3c, Extended Fig. 6b**). As GAD2 is concentrated in GABAergic terminals and its level is regulated by activity^41^, the suppression of *Gad2* translation suggests reduced activity-dependent GABA release at presynaptic terminals of these inhibitory interneurons. Impairment of PV interneuron function could explain the abnormal gamma oscillations observed in *Grin2a*+/- mice^19^ and in human SCZ^42^. Among all brain regions, the inhibitory neurons of the TRN, involved in control of attention, sleep spindles, and sensory filtering^43^, showed the most extensive translational reduction of *Gad2* (**Fig. 3g bottom row**), correlated with the high abundance of PV neurons in this structure. Bulk analysis further revealed translational downregulation of *Gad1* in the SSC (**Supplementary Table 5**), paralleling the GABAergic dysfunction observed in human SCZ studies^37^. As further evidence of disturbed GABAergic signaling in *Grin2a*+/- cortex, we observed translational upregulation of *Slc6a1* (GAT1, mediates GABA reuptake from synaptic cleft) in both TEGLU and TEINH (**Fig. 3a,c**). Beyond the GABAergic system, cortical inhibitory neurons also showed reduced translation of *Sst* (somatostatin), *Cck* (cholecystokinin), and *Gnao1* (G-protein α subunit o) (**Fig. 3a,g**), particularly in the superficial cortical layers, suggesting compromised signaling by Sst and Cck subtypes of inhibitory interneurons and altered G-protein-mediated inhibitory modulation.

Besides reduced translation of key GABAergic-signaling related proteins in TEINH, our spatial translatome profiling revealed additional evidence of disrupted inhibition in glutamatergic neurons of the *Grin2a*+/- brain. In TEGLU, we observed robust translational downregulation (with minimal to slight reduction of mRNA) of *Gabra1*, encoding the α1 subunit of GABAA receptors, suggesting reduced responsiveness to inhibitory inputs to cortical pyramidal neurons (**Fig. 3a**). Translation of *Gabra1* fell also in TEINH neurons (**Fig. 3c**). Of interest, previous studies of human SCZ brain tissue also found reduced expression of GABRA1 in the prefrontal cortex^44^. Notably, we also observed translational downregulation of carbonic anhydrase *Car2* in TEGLU, which may further compromise GABAergic neurotransmission by reducing bicarbonate-dependent modulation of GABAA receptor activity^45,46^. The widespread inhibitory dysfunction, combined with impaired expression of plasticity- and glutamatergic synapse-related genes in TEGLU, suggests a pathological microcircuit state in *Grin2a*+/- cortex characterized by weakened inhibitory control over dysfunctional pyramidal neurons.

MSNs serve as the principal output cells of the basal ganglia, integrating excitatory inputs from cortex and thalamus and dopamine input from substantia nigra and VTA to regulate movement, reward-based learning, decision making, cognitive flexibility and motivation^47^. Direct pathway MSN (MSN_1-[*Drd1*_*Rasd2*]) and indirect pathway MSN (MSN_2-[*Drd2*_*Penk*]) often work in opposition to facilitate or suppress actions, respectively, and disruption of this balance has been consistently implicated in SCZ^48,49^. In our dataset, both MSN subtypes showed translational reduction of *Gad1* and *Gad2*, suggesting reduced GABA synthesis (**Fig. 3d, Supplementary Table 4**). Beyond the shared reduction in GABA synthesis, MSN subtypes exhibited significant translational downregulation of dopamine-receptor signaling pathways (**Fig. 4b**), including both direct pathway (*Drd1*, *Ppp1r1b*) and indirect pathway (*Adora2a*) markers (**Fig. 3d, Supplementary Table 4**). Notably, *Comt*, which catabolizes dopamine, showed translational downregulation in line with the previous report showing downregulation of *Comt* at both mRNA and protein levels^19^. Collectively, these data suggest reduced dopamine catabolism and perhaps heightened dopamine signaling, accompanied by translational downregulation of dopamine receptors in the striatum. Of particular interest, *Chrm1* and *Chrm4*, encoding the M1 and M4 muscarinic receptors and the target of Cobenfy, the recently FDA-approved SCZ drug, were significantly downregulated in MSN_1 and MSN_2, respectively, suggesting disrupted cholinergic signaling in the *Grin2a*+/- striatum (**Fig. 3d, Supplementary Table 4**). Moreover, translational reduction of *Dlg2*, *Dlgap3*, *Nrxn2*, and *Synpo* suggests impact of *Grin2a*+/- on excitatory synapse structure or function in MSNs.

**Fig. 4:**
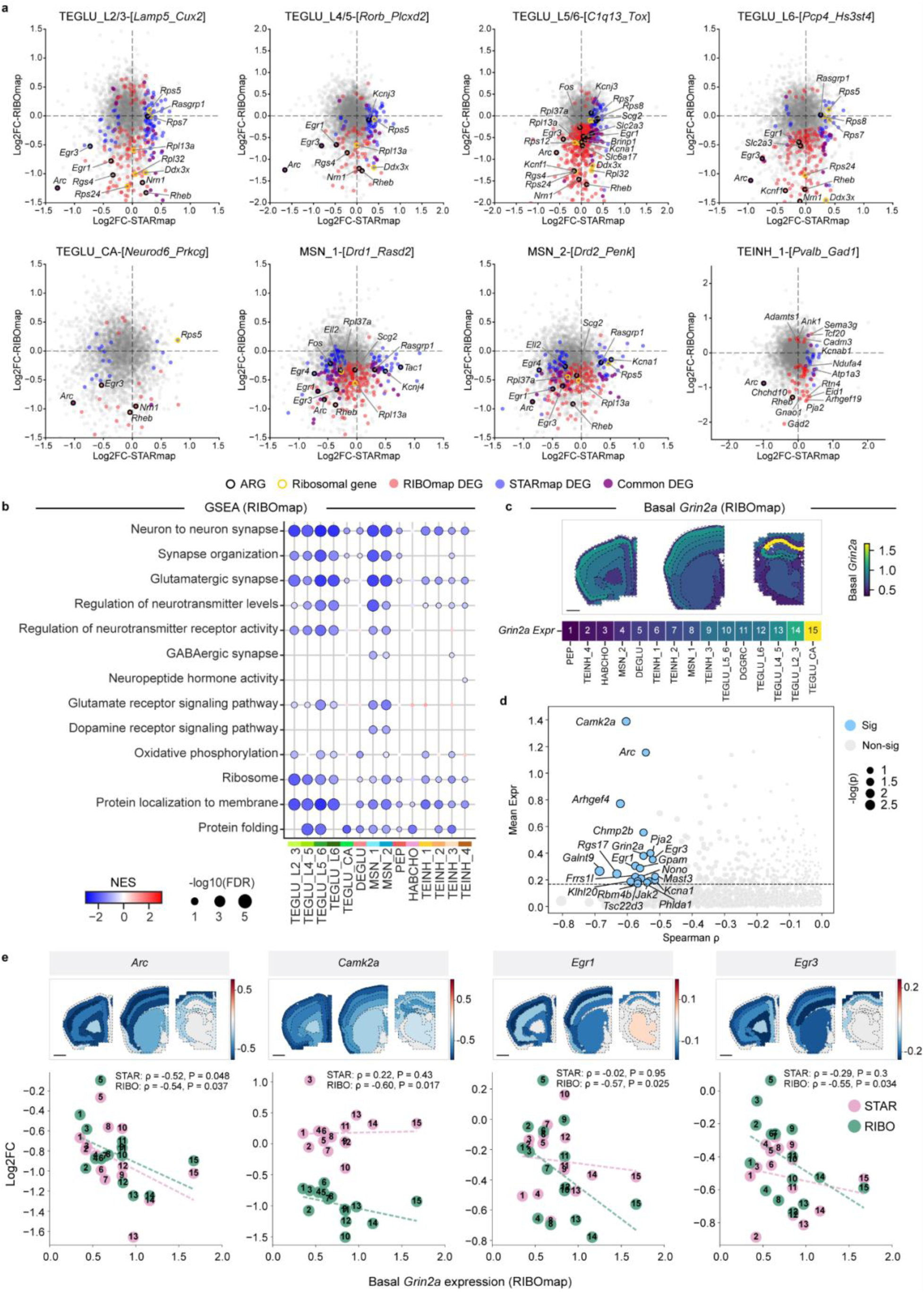
Intrinsic *Grin2a* expression predicts vulnerability to LoF across neuronal subtypes. **a**, STARmap vs. RIBOmap log fold-change plots of neuronal subtypes of interest: TEGLU_L2/3, TEGLU_L4/5, TEGLU_L5/6, TEGLU_L6, TEGLU_CA, MSN_1, MSN_2, and TEINH_1. DEGs with significant (adjusted p-value < 0.05, Log2FC ≥ 0.2) log fold-change in STARmap only, RIBOmap only, and both modalities are colored in blue, red, and purple, respectively. Activity-regulated genes (ARGs) and ribosomal genes identified as DEGs were highlighted with black and yellow circles, respectively. **b**, Gene set enrichment analysis (GSEA) of DEGs identified for neuronal subtypes in RIBOmap. Color scale, normalized enrichment score (NES); dot size, negative log-transformed FDR q-value. **c**, Spatial map of RIBOmap basal *Grin2a* expression (top) and ranking of neuronal subtypes based on RIBOmap basal *Grin2a* expression (bottom). **d**, Scatter plot showing genes with negative Spearman’s ρ between basal *Grin2a* expression and RIBOmap log fold-change across neuronal subtypes. Genes with a p-value lower than 0.05 and passing the 75% expression percentile across neuronal subtypes were labeled. Dot size, negative log-transformed p-value. **e**, Relationship between basal *Grin2a* expression levels and translation reduction of example genes across neuronal subtypes (bottom) and spatially heterogeneous translation reduction of these genes across tissue regions (top).

As RIBOmap identified perturbations of multiple synaptic signaling and plasticity pathways, we next performed quantitative mass-spectrometry (MS) proteomics on synaptic fractions (see **Methods**) purified from the cortex of *Grin2a* LoF mutant mice to assess whether translational alterations (RIBOmap) result in measurable protein-level changes at synapses. To better compare with the synaptic proteome from bulk biochemical preparation, we generated comparable STARmap and RIBOmap profiles combining reads from cell bodies and processes (**Extended Data Fig. 5b-c, Supplementary Table 6**). We observed markedly higher concordance between synaptic proteome profiling and RIBOmap than with STARmap in the *Grin2a*+/- mutant (**Extended Data Fig. 5d,e**), with substantially more shared DEGs. Among the RIBOmap DEGs showing significant changes in the synaptic proteome, 129 out of 197 and 181 out of 255 showed consistent direction in protein-level change in *Grin2a*+/- and *Grin2a*-/- mice, respectively. Most shared DEGs were reduced in both datasets, particularly among postsynaptic signaling, vesicle trafficking, and protein quality-control pathways. Notably, different isoforms of the CAMKII complex (*Camk2a*, *Camk2b*, *Camk2g*) and multiple components of the ESCRT complex (*Chmp3*, *Chmp4*, *Chmp2a*) were downregulated in the synaptic proteome of *Grin2a*+/-or *Grin2a*-/- mutants, as well as in RIBOmap (*Grin2a*+/-), suggesting impaired postsynaptic calcium-dependent signaling and membrane trafficking (**Extended Data Fig. 5d**). Similarly, reduced translation and reduced protein levels of E3 ubiquitin ligases (*Pja1*, *Pja2*) and Hsp40 co-chaperones (*Dnaja2*, *Dnaja4*) suggest compromised proteostasis and synaptic protein turnover. Together, our cross-modal analysis (translatomics and proteomics) revealed that *Grin2a* heterozygosity elicits major translational perturbations across diverse neuronal populations in a cell-type- and brain-region-specific manner, ultimately driving changes in the level of key synaptic proteins whose dysfunction could underlie the circuit aberrations of SCZ.

### Intrinsic *Grin2a* expression predicts vulnerability to LoF across neuronal subtypes

Across neural subtypes, our spatial multi-omics approach revealed shared changes upon *Grin2a* loss, particularly in RIBOmap (**Fig. 4a, Extended Data Fig. 6a-c**). A prominent pattern of translational downregulation was observed for canonical activity-regulated genes^50^ (ARGs) (*Egr1*, *Egr3*, *Arc*, *Rheb*, *Nrn1*), ribosomal proteins (*Rpl13a*, *Rps24*, *Rpl32*), cytoplasmic signaling proteins (*Rasgrp1*, *Rgs4*), and transcription/translation/metabolism regulators (*Ddx3x*, *Ell2*, *Slc2a3*, *Scg2*) across multiple TEGLU and MSN subtypes (**Fig. 4a**). GSEA of RIBOmap data confirmed broad translational downregulation of shared pathways not significantly altered at the mRNA level (**Fig. 4b**). GO terms related to synapses, glutamatergic synaptic signaling, translation, protein folding and transport, and oxidative phosphorylation were consistently translationally downregulated across most neuronal subtypes. Spatial mapping of gene set score differences further revealed that activity-related gene sets, ARGs and immediate early genes (IEGs), were broadly reduced in RIBOmap across cortex and hippocampus, most prominently in superficial cortical layers (**Extended Data Fig. 6d**). Meanwhile, oxidative phosphorylation and ribosomal programs showed opposite regulation in RIBOmap in cortical regions versus striatum, despite their mild upregulation in STARmap across most regions (**Extended Data Fig. 6d**), suggesting region-specific translation control. These findings indicate that *Grin2a* loss disrupts translation of a common set of genes and pathways critical for synaptic function and neuronal activity, with magnitude varying across neuronal subtypes.

We next hypothesize that the magnitude of translational changes reflects the varying dependencies of neuronal subtypes on *Grin2a* for maintaining their functional state. To test this, we sought to correlate basal *Grin2a* expression in wild-type cells with RIBOmap log fold-changes of individual DEGs across 15 neuronal subtypes. It was worth noting that TEGLU subtypes had higher baseline *Grin2a* expression than other neuronal populations, with TEGLU_CA showing the highest levels, while cortical TEGLU subtypes displayed a decreasing gradient of *Grin2a* expression from superficial to deep layers (**Fig. 4c**). Next, we applied the Spearman’s rank correlation test to identify genes with RIBOmap log fold-change highly correlated with basal *Grin2a* expression across neuronal subtypes (**Fig. 4d, Supplementary Table 7**). Our analysis revealed a set of *Grin2a*-dependent genes with pathway-level associations, consisting of key ARGs and synaptic plasticity regulators such as *Arc*, *Camk2a*, *Arhgef4*, *Chmp2b*, *Egr1*, *Egr3*, and *Pja2* (**Fig. 4d, Extended Data Fig. 6e**). Specifically, we observed a quantitative relationship where neuronal subtypes with higher basal *Grin2a* expression experienced greater translational repression of these genes (**Fig. 4e**). From a spatial perspective, this subtype heterogeneity indicated higher vulnerability of cortical layers and the hippocampus compared to subcortical regions, aligning with gsMap (genetically informed spatial mapping of cells for complex traits) prediction based on human genomics data^51^ (**Extended Data Fig. 7**). In contrast, the correlation between *Grin2a* basal expression and differential expression was weaker or absent in STARmap data (**Fig. 4e**), particularly for *Camk2a*, *Egr1*, and *Egr3*, suggesting that the gene dosage effect is primarily mediated at the translational level. *Arc*, however, showed significant correlation between STARmap log fold-change and basal *Grin2a* expression, suggesting predominantly transcriptional regulation. This set of *Grin2a*-dependent genes quantitatively links *Grin2a* dosage to translational changes of downstream genes, partially explaining the heterogeneous effects of *Grin2a* LoF across neuronal types.

### Transcriptional and translational changes in non-neuronal cell types of *Grin2a*+/- mice

Glial and vascular cells also showed RIBOmap and STARmap changes in *Grin2a+/-* mutants (**Fig. 5a-d, Extended Data Fig. 8a**) presumably secondary to at least some neuronal alterations, as *Grin2a* is expressed at low to minimal levels in non-neuronal cells (**Extended Data Fig.5a**). Astrocytes showed translational reduction of GABA transporters *Slc6a11* and glutamate transporter *Slc1a3*, suggesting altered clearance of inhibitory and excitatory neurotransmitters (**Fig. 5a**). Key regulators of ionic and osmotic homeostasis, *Aqp4* and *Car2*, were downshifted at the translational level, suggesting impaired water flux and pH buffering at astrocytic endfeet^52,53^. Moreover, genes involved in lipid/cholesterol synthesis (*Hmgcs1*, *Plpp3*, *Gpam*) and *Apoe*, were downregulated in RIBOmap, suggesting translation reduction of astrocytic lipid/cholesterol production and export. Notably, *Insig1*, a negative regulator of cholesterol synthesis, was upregulated at the translational level, in line with mRNA upregulation of this gene observed in previous snRNA-seq study^19^. GSEA results further revealed translational downregulation of lipid/cholesterol metabolism, along with glutamate secretion and chaperone binding, in astrocytes (**Fig. 5e**). Given that astrocytes are the major source of neuronal cholesterol, disrupted astrocytic lipid supply could impair synaptic stability/function and membrane turnover in neurons^54,55^. Beyond this, we also observed altered translation of astrocytic genes involved in blood-brain barrier function, including *Gja1*, *Hepacam*, *Vegfa*, and *Sema3g* (**Fig. 5a**).

**Fig. 5:**
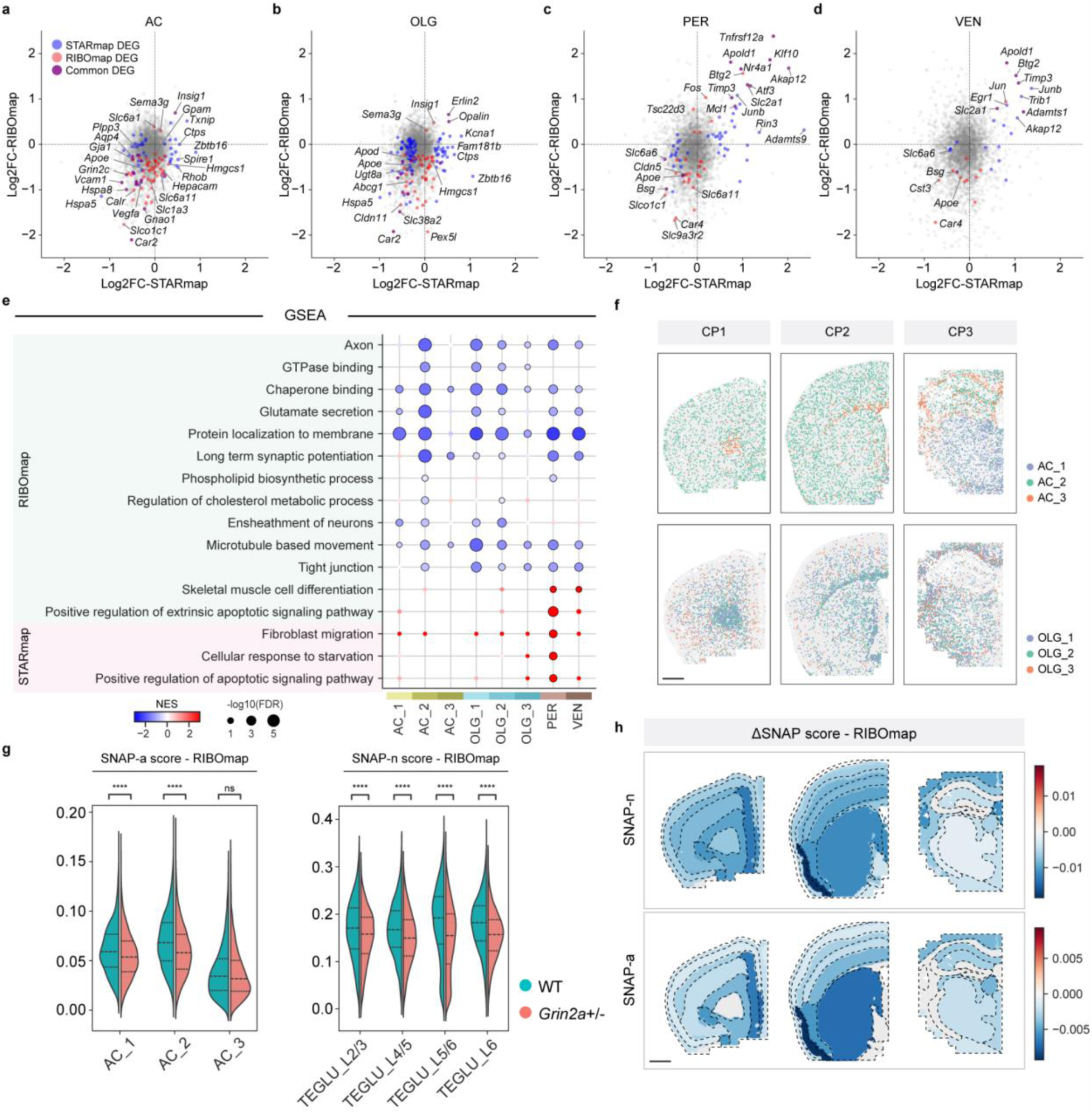
Transcriptional and translational changes in non-neuronal cell types. **a-d**, STARmap vs. RIBOmap log fold-change plots of astrocytes (AC), oligodendrocytes (OLG), pericytes (PER), and vascular endothelial cells (VEN). DEGs with significant (adjusted p-value < 0.05, |Log2FC| ≥ 0.2) log fold-change in STARmap only, RIBOmap only, and both modalities are colored in blue, red, and purple, respectively. **e**, GSEA of DEGs identified in astrocytes, oligodendrocytes, pericytes, and vascular endothelial cells. Color scale, normalized enrichment score (NES); dot size, negative log-transformed FDR q-value. **f**, Spatial visualizations showing the subtype distribution of astrocyte (top) and oligodendrocyte (bottom) at 3 coronal positions. Scale bar, 1 mm. **g**, Violin plots illustrating the difference between WT and *Grin2a*+/- in SNAP-a gene program scores in astrocyte subtypes (left) and SNAP-n gene program scores in cortical TEGLU subtypes (right) in RIBOmap. The P value was calculated using a two-sided independent t-test. ****P < 0.0001. **h**, Spatial visualizations of SNAP-n score difference in TEGLU cortical neurons (top) and SNAP-a score difference in astrocytes (bottom) in RIBOmap. Scale bar, 1 mm.

In oligodendrocytes, RIBOmap also identified downregulation of genes involved in lipid/cholesterol synthesis and transport (*Hmgcs1*, *Ugt8a*, *Apoe*, *Apod*, *Abcg1*), accompanied by upregulation of *Insig1* and *Erlin2* (involved in cholesterol metabolism and ER stress response) (**Fig 5b**). Lipids are important for myelin formation and integrity, as are tight junction protein *Cldn11* and carbonic anhydrase *Car2*, which were also downregulated in RIBOmap, suggesting compromised myelination in *Grin2a+/-* mutant brain^56^. In contrast, *Opalin*, an oligodendrocyte-specific glycoprotein enriched at paranodal and inner myelin loops, was upregulated at both mRNA and translational levels (**Fig. 5b**). Oligodendrocyte precursor cells (OPC) showed translational downregulation of *Pdgfra*, a canonical marker and key regulator of OPC proliferation (**Extended Data Fig. 8a**). Compared with astrocytes and oligodendrocytes, microglia showed fewer translational changes, but purinergic receptor *P2ry12*, G protein-coupled receptor *Gpr34*, fractalkine receptor *Cx3cr1*, and lysosomal protease *Ctss*, were downregulated in RIBOmap^57^ (**Extended Data Fig. 8a**).

Vascular cell populations, primarily pericytes and vascular endothelial cells, exhibited strong concordance between transcriptional and translational changes for upregulated DEGs (**Fig. 5c,d**), including glucose transporter *Slc2a1*, angiogenesis regulator *Apold1*, and extracellular-matrix regulators *Adamts1* and *Timp3*. Interestingly, IEGs (*Btg2*, *Jun*, *Junb*, *Klf10*, *Nr4a1*) and stress-responsive regulators^58–60^ (*Tnfrsf12a*, *Akap12*, *Tsc22d3*) were elevated at both mRNA and translational levels. Among the translationally downregulated genes, carbonic anhydrase *Car4* and tight junction protein *Cldn5* stood out, suggesting perturbation of perivascular pH homeostasis and blood-brain barrier integrity. Moreover, GSEA suggested translational downregulation of the tight-junction pathway in vascular cells (**Fig. 5e**). At the transcriptional level, pericytes displayed the strongest signals associated with apoptosis, fibroblast migration, and nutrient-stress pathways (**Fig. 5e**). Together, these findings suggest vascular stress and altered blood brain barrier function in *Grin2a+/-* brain.

Additionally, we observed significant differences in DEG profiles among subtypes of astrocytes and oligodendrocytes (**Extended Data Fig. 8b,c**), each characterized by unique marker genes and distinct spatial distributions (**Fig. 5f**). Specifically, AC_2-[*Cspg5*_*Mfge8*], located within cortico-striatal and hippocampal regions (where neurons showed translational changes in numerous genes including ARGs) showed the highest number of DEGs among astrocyte subtypes (**Extended Data Fig. 8b**). In contrast, white-matter localized AC_3-[*Gfap*_*Aqp4*], with little local contact with neurons, showed significantly fewer DEGs at both mRNA and translational levels. Similarly, the white-matter-localized oligodendrocyte subtype (OLG_3) showed the least transcriptional and translational changes among all three subtypes (**Extended Data Fig. 8c**). These data further support the notion that, in *Grin2a* +/- animals, glial alterations are secondary to initiating changes in local neurons, where *Grin2a* is primarily expressed. Analyses by snRNA-seq of human postmortem prefrontal cortex have revealed concerted regulation of a synaptic neuron and astrocyte program (SNAP), which is under-expressed in SCZ patients^61^; we calculated SNAP scores (using the mouse orthologs of these human genes) for astrocytes (SNAP-a) and neurons (SNAP-n) to test whether such a concerted downregulation also occurs in *Grin2a+/-* mouse. Indeed, all cortical TEGLU subtypes showed significant translational reduction of the SNAP-n program, consistent with the idea that decline in SNAP disrupts the synaptic proteome; we also observed significant translational reduction of the SNAP-a program in AC_1 and AC_2 (grey matter astrocytes), but not in AC_3 (white matter astrocytes) (**Fig. 5g**). Furthermore, the RIBOmap-measured changes of SNAP-a and SNAP-n showed striking similarities in their spatial distributions, consistent with the idea that they are coordinated (**Fig. 5h**). Both SNAP-a and SNAP-n exhibited the greatest reduction in the insular cortex (CTX_AI2/3), a brain region responsible for processing sensory stimuli and highly affected in SCZ^62^, and were more downregulated in the prefrontal cortex and the striatum compared to the somatosensory cortex and thalamus. In comparison, STARmap captured little evidence of astrocyte-neuron coordination: SNAP-a was decreased in all astrocyte subtypes while SNAP-n was significantly changed only in TEGLU_L4/5 (upregulated) (**Extended Data Fig. 8d,e**). Thus, both cell-subtype and spatial analyses suggest coordinated regulation between neuronal synaptic function and local astrocyte support manifests at translational level and is undermined by *Grin2a*+/-.

### Co-variation of translatomic changes among neurons and non-neuronal cell types across brain regions in *Grin2a*+/- mice

Leveraging the spatial resolution of RIBOmap data, we next examined whether different brain cell types within the same tissue region have coordinated translatome changes. To this end, we first computed region-specific RIBOmap log fold-changes of selected DEGs from four abundant cell populations, including neurons (169 DEGs shared between at least 3 neuronal subtypes), astrocytes (64 DEGs), oligodendrocytes (56 DEGs), and pericytes (39 DEGs) (**Supplementary Table 8**). Five molecular tissue regions (VS, FT, MNG, CTX_HIP, CTX_PIR) were excluded from the analysis due to low neuron abundance, leaving 18 regions for downstream correlation analysis. We computed Pearson’s correlation coefficients for all possible cross-cell-type DEG pairs across these regions as a metric of gene-gene co-variation. Our analysis revealed significant co-variations between neuronal and non-neuronal DEGs (**Fig. 6a**): 219 neuron-astrocyte pairs, 56 neuron-oligodendrocyte pairs, and 39 neuron-pericyte pairs exhibited strong co-variation (|Pearson’s r|>0.8). In addition, we identified 22 highly correlated DEG pairs among different non-neuronal cell types: 10 between astrocytes and oligodendrocytes, 7 between astrocytes and pericytes, and 5 between oligodendrocytes and pericytes (**Fig. 6a**).

**Fig. 6:**
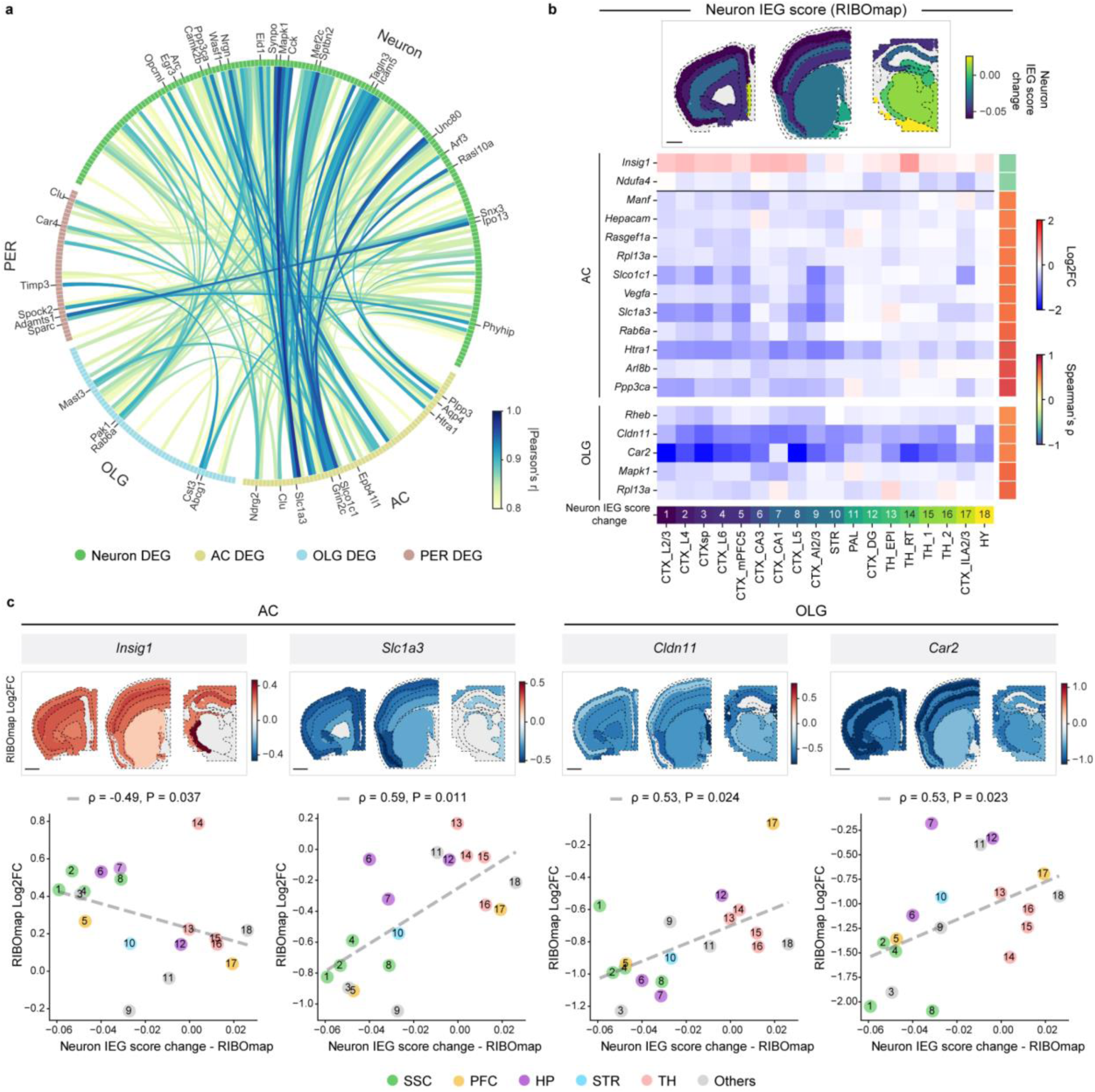
Translational co-variation among neuronal and non-neuronal cell types across brain regions. **a**, Chord diagram showing pairwise correlations of DEGs across neuron, astrocyte, oligodendrocyte, and pericyte based on region-specific RIBOmap log fold-change. Chord color, absolute value of pairwise Pearson’s correlation coefficient r. **b**, Heatmap showing significant correlations between the regional log fold-change of glial DEGs from RIBOmap and changes in regional neuron RIBOmap IEG scores (bottom, p-value < 0.05), along with the spatial distribution of these IEG score changes (top). **c**, Relationship between regional neuron RIBOmap IEG score change and RIBOmap log fold-change of example glial DEGs (bottom), spatial heterogeneity of RIBOmap log fold-change of these genes across tissue regions (top).

Notably, astrocytes exhibited the most extensive gene-gene translational co-variations with neurons (**Fig. 6a**), consistent with their role in supporting and responding to synapses^63,64^. Given that the neurovascular unit (NVU) relies on interplay between neurons, astrocytes, and vascular cells^65^, we next examined whether cross-cell-type co-variations exist among NVU-related genes and indeed found a dense interaction network linking neuronal, astrocytic, and pericytic NVU-associated DEGs (**Extended Data Fig. 9a,b**). In particular, the astrocytic glutamate transporter *Slc1a3* (encoding EAAT1) showed high correlation with vesicular glutamate transporter *Slc17a7* and multiple genes in NMDAR-dependent signaling pathways (*Camk1*, *Ppp3ca*, *Arc*) (**Extended Data Fig. 9a**), suggesting that regional changes in astrocytic glutamate clearance are coordinated with local glutamatergic signaling-associated changes in neurons. Other representative co-varied DEG pairs included *Camk1* (neuron) with *Slc1a3* (astrocyte), *Egr3* (neuron) with *Htra1* (astrocyte), *Atp1a1* (neuron) with *Spock2* (pericyte), and *Vegfa* (astrocyte) with *Timp3* (pericyte) (**Extended Data Fig. 9b**). These patterns suggest that translational shifts in neurons, astrocytes, and pericytes are interlocked, coordinating neuron state with glial and vascular support.

To pinpoint glial DEGs with direct relevance to local neuronal activity-associated gene programs, we first derived a “neuronal activity index” for each region by calculating the mean gene group score of IEGs by RIBOmap in neurons. We then correlated the regional change in neuronal IEG score with the log fold-change of every glial DEG, using Spearman’s rank correlation to capture monotonic relationships (**Fig. 6b**). This analysis revealed a set of glial RIBOmap DEGs (13 astrocyte DEGs, 5 oligodendrocyte DEGs) that appear coupled to neuronal activity, such as *Insig1*, *Slco1c1*, *Vegfa*, *Slc1a3*, and *Htra1* in astrocytes, and *Cldn11*, *Car2* in oligodendrocytes. Interestingly, *Slco1c1* and *Vegfa* were previously identified as astrocytic activity-response genes (AAR genes) in neuron-astrocyte co-culture systems, which corroborates our discovery of their gene-activity co-variation in brain tissues^66^. In cortical areas, where neuronal activity-associated programs show prominent translational changes, these glial activity-coupled genes tend to show greater differential expression (**Fig. 6c,d**). For example, astrocytes suppress *Slco1c1*, *Slc1a3*, and upregulate *Insig1*, while oligodendrocytes downregulate *Car2* (**Fig. 6d**). This concerted modulation implies a stronger cortical disruption of lipid metabolism, neurotransmitter clearance, and pH-buffering. In contrast, in thalamic regions where neuronal activity-associated programs showed subtle changes, modest differential expressions were observed for the same genes. Notably, the neuron-activity-coupled astrocytic translation shift of *Insig1* paralleled human schizophrenia snRNA-seq findings showing downregulation of astrocytic cholesterol biosynthetic genes in correlation with reduced neuronal activity markers^61^. Together, these findings indicate that glial support programs are not uniformly changed across the brain in *Grin2a*+/- mutants but instead scale translationally with local neuron state changes as measured by RIBOmap.

## Discussion

By integrating spatially resolved transcriptomic and translatomic technologies, we revealed pronounced translational dysregulation across the *Grin2a*+/- mutant brain with striking cell-type and regional heterogeneity. In fact, numerous RIBOmap DEGs showed marked changes in translation but little change in mRNA level (*Ppp3ca*, *Gabra1*, *Gad2*, *Rheb*, *Camk2a*, *Nrn1* being interesting examples). Our findings underscore the critical role of translational control in brain function and SCZ pathophysiology and highlight an important regulatory mechanism that is quite overlooked in transcriptome-only studies.

Our study uncovered translational dysregulation of excitatory and inhibitory neurotransmission in major neuron cell types, including glutamatergic, GABAergic, and striatal dopaminergic signaling. Moreover, we identified pervasive translational suppression of calcium-dependent synaptic plasticity regulators across multiple neuronal subtypes, including *Arc*, *Egr1*, *Rheb*, *Camk2a*, and *Ppp3ca*, which may contribute to SCZ-associated deficits in learning and memory processes. By comparing RIBOmap with synaptic proteomics, we identified a set of RIBOmap DEGs and synaptic DEPs that change in the same direction and are known to act in synaptic function and plasticity, including CAMKII isoforms, ESCRT-associated endosomal sorting factors, E3 ubiquitin ligases, and Hsp40 co-chaperones. Our results suggest that the compromised protein levels of these genes at synapses is driven by translational suppression of their mRNAs, rather than by reduced mRNA levels.

RIBOmap was also able to uncover prominent alterations in inhibitory interneurons (TEINH) that were absent in STARmap and that were largely missed by previous snRNAseq analysis of *Grin2a*+/- brain^19^. Notably, reduced translation of GABA synthesizing enzyme *Gad2* in different TEINH subtypes, most prominently in PV interneurons (**Extended Data Fig. 6b**), as well as downregulation of neuropeptides cholecystokinin (*Cck*) and somatostatin (*Sst*) which are primarily expressed by inhibitory interneurons (**Fig. 3c**) suggests disrupted inhibitory interneuron function. Our findings in TEINH are reminiscent of human postmortem SCZ studies, which also uncovered reduced expression of GAD in PV interneurons of prefrontal cortex^36,38,39^, and are in keeping with a long-standing theory for the cognitive impairment of SCZ^37^. Thus, together with our prior transcriptomic study^19^, the current data show evidence in support of the three major hypotheses for the pathophysiology of SCZ in *Grin2a*+/- mice: hypoglutamatergic function^15,16^; hyperdopaminergic signaling in the striatum^67^; and inhibitory interneuron dysfunction in the neocortex^37,38^.

By correlating baseline *Grin2a* expression with translational changes across 15 neuronal subtypes, we provided key molecular insights into cell autonomous impacts of *Grin2a* LoF on downstream genes. We identified a set of genes, including *Camk2a*, *Arc*, *Egr1*, *Egr3*, *Chmp2b*, and *Pja2*, whose translational suppression was highly correlated with basal (WT) expression of *Grin2a* across neuronal subtypes, with this dosage effect operating primarily translationally rather than transcriptionally. Cortical and hippocampal TEGLU with the highest basal *Grin2a* expression exhibited the greatest translational dysregulation of these genes, which aligns with these cell - types’ particularly strong association with human SCZ traits^51^ (**Fig. 4c,d**, **Extended Data Fig. 7**), demonstrating high human SCZ-relevance of our finding.

Beyond neurons, our study revealed distinct and region-specific translational alterations in non-neuronal cell types. In RIBOmap, astrocytes showed translational downregulation of neurotransmitter transporters and lipid/cholesterol metabolism genes, indicating compromised synaptic support, particularly in cortical regions. Oligodendrocytes exhibited dysregulated lipid metabolism and myelin integrity maintenance. It is noteworthy that no significant STARmap changes were observed for most RIBOmap DEGs in the aforementioned pathways, suggesting RIBOmap (translatomic profiling) is more capable of detecting differential expression indicative of functional changes also in glial cell types. Meanwhile, vascular cells upregulated stress-response and extracellular matrix remodeling programs. Given that these non-neuronal cell types express *Grin2a* at low to minimal levels, we reason that the translatomic changes in these cells are secondary to changes in neurons.

The spatial resolution of STARmap and RIBOmap allows us to uncover regional heterogeneity in cell-cell interactions that would have been missed by bulk approaches. Across 18 molecular tissue regions, we revealed striking cross-cell-type DEG co-variations between neurons, astrocytes, oligodendrocytes, and pericytes, particularly among NVU-associated genes. Additionally, we discovered astrocyte and oligodendrocyte DEGs that co-varied with RIBOmap IEG score in neurons in the local region. These findings highlight a coordinated local response to NMDAR hypofunction, where glial support programs scale with local neuronal circuit disturbance.

Furthermore, our study bridges the findings in *Grin2a*+/- mouse model with previous characterization of human SCZ patients. At subcellular level, we identified key RIBOmap DEGs, such as *Grin1* and *Homer1*, that also show significant reduction in synaptic proteome of human SCZ samples^68,69^. At cell-cell interaction level, we observed a concerted disruption of the neuron-astrocyte program SNAP, a gene set showing concerted downregulation in postmortem prefrontal cortex of SCZ patients. At the brain-region level, we identified prominently affected regions in *Grin2a*+/- model, such as superficial cortical layers, prefrontal cortex, insular cortex, and thalamic reticular nucleus, which aligns with computational predictions based on human genetics data (**Extended Data Fig. 7b**). These consistencies largely reinforce the disease relevance of our findings in the *Grin2a*+/- model.

In summary, our integrated spatial omics analysis shows that *Grin2a* haploinsufficiency drives brain-wide translational shifts in all cell types examined. Translational reduction in calcium-dependent synaptic plasticity and neurotransmission appears to be a widespread consequence of NMDAR hypofunction in both excitatory and inhibitory neurons, along with further cell-type and region-specific adaptations across neurons, glia, and vascular cells. Given the prominent translational dysregulation of *Grin2a*+/- model, our study inspires future applications of RIBOmap to explore the spatial molecular changes in other SCZ disease models and human patient samples.

## Methods

### Mouse brain tissue collection

All animal procedures followed animal care guidelines approved by the Institutional Animal Care and Use Committee (IACUC) at the Broad Institute of MIT and Harvard. Male *Grin2a*+/- and wild-type littermate mice were anesthetized with isoflurane, perfused with PBS, and decapitated at 12 weeks of age. Brains were rapidly extracted, snap freezed in O.C.T. with liquid nitrogen, and stored at −80 °C.

### STARmap and RIBOmap

We employed paired primer and padlock probes in STARmap to detect mRNA irrespective of its translational state, whereas a combination of splint, primer, and padlock probes was used in RIBOmap to specifically identify ribosome-bound mRNA^5^.

In detail, mouse brain samples were sectioned at 20 µm thickness on Leica Cm1950 cryostat at −20 °C. For each animal, 6 coronal sections, including 2 adjacent sections at each of the 3 coronal positions, were prepared and transferred to 24-well glass-bottom plates (Cellvis). Plates were pre-treated by oxygen plasma cleaning, followed by bind-silane coating (10% acetic acid, 1% methacryloxypropyltrimethoxysilane, 89% ethanol; 1 h, room temperature). Plates were rinsed three times with 95% ethanol, air-dried, and coated with 50 µg/mL poly-D-lysine (1 h, room temperature). Tissue sections were fixed in 4% paraformaldehyde (PFA) in PBS (15 min, room temperature) and permeabilized in pre-chilled methanol (–20 °C, 1 h). Sections were then quenched in PBSTR buffer (0.1% Tween-20, 0.1 U/µL RNase inhibitor in PBS) supplemented with 1% yeast tRNA and 100 mM glycine (5 min, room temperature), followed by rinsing in PBSTR.

For STARmap and RIBOmap hybridization, probe mixtures were prepared by combining equal volumes of diluted probe solution (padlock and primer probes, 2 nM per oligo) with hybridization buffer (4x SSC, 20% formamide, 2% Tween-20, 40 mM ribonucleoside vanadyl complex, 0.2 mg/mL yeast tRNA, 0.4 U/µL SUPERase inhibitor), yielding a final probe concentration of 1 nM per oligo. For RIBOmap, an additional splint probe (100 nM final concentration) was included. Sections were incubated in a hybridization mixture at 40 °C in a humidified chamber with gentle shaking and parafilm sealing for 36 h. Excess probes were removed by two PBSTR washes followed by a high-salt wash (4x SSC in PBSTR) at 37 °C for 20 min each.

Padlock ligation was performed in T4 DNA ligase buffer containing 0.5 mg/mL BSA, 0.2 U/µL RNase inhibitor, and 0.25 U/µL T4 DNA ligase (3 h, room temperature), followed by 3 PBSTR washes. Following ligation, sections were incubated in rolling circle amplification (RCA) solution consisting of 0.5 U/µL Phi29 DNA polymerase, 250 µM dNTPs, 20 µM 5-(3-aminoallyl)-dUTP, 0.2 mg/mL BSA, and 0.2 U/µL RNase inhibitor in Phi29 buffer. Samples were preincubated at 4 °C for 30 min, then transferred to 30 °C for 2 h with gentle shaking, and subsequently washed twice in PBST (0.1% Tween-20 in PBS).

For hydrogel embedding, sections were treated with modification solution (25 mM methylacrylic acid NHS ester in 100 mM sodium bicarbonate) for 1 h at room temperature, rinsed twice in PBST, and incubated in monomer buffer (4% acrylamide, 0.2% bis-acrylamide, 2x SSC, 0.2% TEMED) for 10 min at 4 °C. The buffer was removed, and 40 µL of polymerization mixture (0.2% ammonium persulfate in monomer buffer) was applied to each section. Gel Slick-coated coverslips were placed immediately over the samples, and polymerization proceeded for 1.5 h at room temperature under a nitrogen atmosphere. Coverslips were carefully removed, and hydrogels were washed twice in PBST. Sections were digested in proteinase K solution (0.2 mg/mL proteinase K, 1% SDS, 2x SSC) at 37 °C for 1 h, followed by PBST washes. Dephosphorylation was performed in Antarctic phosphatase buffer containing 0.25 U/µL Antarctic phosphatase and 0.2 mg/mL BSA at 37 °C for 1 h, followed by PBST washes.

Each SEDAL cycle began with two 10 min treatments at room temperature in stripping buffer (60% formamide, 0.1% Triton X-100), followed by three 5 min PBST washes. Samples were then incubated for 3 h at room temperature in sequencing buffer (1x T4 DNA ligase buffer, 0.2 mg/mL BSA, 10 µM reading probe, 5 µM fluorescent decoding probe, 0.2 U/µL T4 DNA ligase). After incubation, samples were washed three times with imaging buffer (2x SSC, 10% formamide) and maintained in the same buffer during imaging. DAPI staining was performed prior to the first sequencing cycle for 3 h.

Imaging was carried out on a Leica TCS SP8 confocal microscope equipped with a 40x oil-immersion objective. For each cycle, signals were acquired in Alexa Fluor 488, 546, 594, and 647 channels. DAPI signals were collected in the first cycle using a 405 nm laser. A total of 8 sequencing-imaging cycles were conducted, enabling the detection of 3,447 target genes.

### 5’ phosphorylated padlock library preparation

The padlock probe library was synthesized by IDT oPool service without 5’ phosphorylation. A common annealing site was added to the 5’ end of all padlock probe sequences (STARmap padlocks: TAATACGACTCACTATACTGCTAGCGACGGCCA; RIBOmap padlocks: TAATACGACTCACTATACTGCTAGATAACACGGCCT). The probe pool was then cleaved by a 13PD-1 DNAzyme cutter oligo (STARmap: AAAAAAAAAAAATGGCCGTCGCTTATACCGGGCAACTATTGCCTCGTCATCGCTATTTTCTGCGATAGTGAGTCGTATTAAAAAAAAAAAAA, RIBOmap: AAAAAAAAAAAAAGGCCGTGTTATCTTATACCGGGCAACTATTGCCTCGTCATCGCTATTTTCTGCGATAGTGAGTCGTATTAAAAAAAAAAAAA) at a final concentration of 100 nM in the annealing buffer containing 50 mM HEPES (pH 7.0 at 22 °C), 100 mM NaCl, and 10 mM MgCl_2_. The mixture was incubated on a thermocycler at 90 °C for 3 min and cooled to 25 °C at a rate of −0.1 °C/s, and then an equal volume of the cutting buffer (50 mM pH 7.0 HEPES, 100 mM NaCl, 10 mM MgCl_2_, 10 mM MnCl_2_, and 4 mM ZnCl_2_) was added. The reaction was then incubated at 37 °C to allow DNAzyme-assisted autocleavage to yield a 5’ phosphate moiety on the padlock probes. The reaction was then quenched by 0.5 M EDTA. The oligos were ethanol precipitated, and the 5’ phosphorylated padlock pools were isolated on an Agilent 1260 Infinity II HPLC with acetonitrile/hexylamine/acetic acid (pH 7.0) mobile phase and PLRP-S stationary phase. The purified 5’ phosphorylated padlocks were pooled, desalted, ethanol precipitated, and resuspended in 0.1 TE buffer, and the concentration of probes was quantified by Qubit ssDNA assay.

### Imaging data processing

Image deconvolution was achieved with Huygens Essential version 24.04 (Scientific Volume Imaging, The Netherlands, http://svi.nl), using the CMLE algorithm, with SNR:10 and 10 iterations. Image registration, spot calling, and barcode filtering were performed using established software Starfinder (https://github.com/wanglab-broad/starfinder) with customized configuration.

### Quality control and preprocessing

After quantifying signals at the single-cell level, we excluded low-quality cells from each replicate based on the number of transcripts and genes per cell. We used the median absolute deviation (MAD) to establish filtering thresholds for reads per cell:

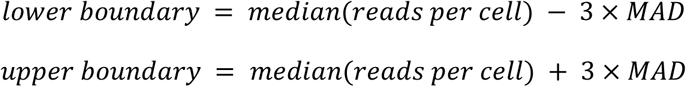

Additionally, we applied standard filtering criteria: 1) A minimum of 10 expressed genes is required for a cell to pass filtering. 2) A minimum of 10 expressing cells is required for a gene to pass filtering. Following these filters, we retained a total of 538,188 cells with 3,447 genes. The expression profiles were normalized using the pp.normalize_total function (Scanpy v1.9.2), followed by logarithmic transformation with pp.log1p. We scaled the data matrix to unit variance using pp.scale and mitigated variations in total reads per cell with pp.regress_out. Finally, to ensure a high-quality cell-typing, we selected 1,017 canonical cell type markers from previous reports to perform a Harmony integration on the processed data matrix to align transcriptome and translatome modalities before conducting downstream analyses such as dimensionality reduction and unsupervised clustering.

### Cell type classification

We applied a hierarchical clustering strategy to create a three-level cell-type annotation for the integrated dataset. First, we constructed a k-nearest neighbor (k-NN) graph from the integrated PCA matrix, connecting cells based on their expression similarity in high-dimensional space. We then used the Leiden community detection algorithm on this k-NN graph to identify cell clusters, resulting in twenty-eight clusters at a clustering resolution of 2. Each cluster was classified as either neuron or glial cell based on canonical markers (e.g., *Slc17a7*, *Gad1*, *Gad2*, *Plp1*, *Slc1a3*). Subsequently, we generated second-level annotations reflecting major cell types such as telencephalon projecting neurons, oligodendrocytes, astrocytes, and vascular cells.

To identify subpopulations of major cell types, we applied the same analyses to various populations under second-level annotation (e.g., telencephalon projecting neurons and interneurons). We used the elbow method to determine significant principal components after performing PCA. By plotting the variance ratio of each principal component with the pl.pca_variance_ratio function, we selected the top 10 to 30 components with the highest values for constructing a kNN graph for clustering. Specifically, eleven clusters were first identified from the telencephalon projecting neurons (TEGLU). The subpopulations located in the hippocampal and cortical regions were then subject to unsupervised clustering again to differentiate neuronal subtypes with distinct spatial representation in the brain regions (i.e., anatomical regions such as L2/3, L4/5, CA, DG, etc.), resulting in seven subpopulations with unique representative gene markers. For inhibitory interneurons, we resolved four clusters distinguished by different neuronal peptide gene markers (e.g., *Pvalb*, *Sst,* and *Vip*). In non-neuronal cells, we identified three subpopulations each from oligodendrocytes and astrocytes, along with five from vascular cells. Additionally, cells without conventional maker expression were labeled as mixed cells and excluded from downstream cell-type-specific analyses.

### Molecular tissue region identification

Tissue region annotation was performed using the SPIN algorithm, which combines single-cell gene expression similarity and spatial proximity to identify tissue regions. Gene expression features for each cell were averaged from a subsampled set of its spatial neighbors. These smoothed features were then clustered using the Leiden algorithm at a resolution of 0.7, producing distinct molecular regions, where most identified regions had representative gene markers. Anatomical labels were then assigned by overlaying these regions with the Allen Mouse Brain Atlas and comparing with the Spatial Mouse Brain Atlas in Shi et al. Finally, the annotated cells were further refined using kNN majority voting (n_neighbors=50) to create clearer tissue boundaries.

### Consensus non-negative matrix factorization (cNMF)

cNMF (v.1.7) was applied to the integrated dataset following the PBMC tutorial on GitHub (https://github.com/dylkot/cNMF/blob/master/Tutorials/analyze_pbmc_example_data.ipynb). Factorization was performed on filtered raw counts data, iterating over k values from 30 to 40. Based on the PCA of the gene expression matrix and the cNMF stability report, we selected k = 35 with a density threshold of 0.2 for further analysis. Additionally, the latent factor usage matrix (cell by factor) was normalized so that each cell’s total usage across all factors summed to one before analysis. With each module, top 50 genes were subjected to downstream analysis.

### Differential gene expression analysis

Differential expression (DE) analysis was conducted using the sc.tl.rank_genes_groups function in Scanpy, employing the Wilcoxon rank-sum test. Comparisons were made between genotypes within each cell type, retaining only genes with an absolute log fold-change (log₂FC) of 0.2 or greater and a Benjamini-Hochberg-adjusted p-value lower than 0.05. To ensure consistency across biological replicates, we applied a filter that included only genes exhibiting changes in the same direction. To ensure cell type specificity, we further applied a filter based on cNMF gene modules to exclude glial DEGs that are potentially dominated by mis-segmented reads from neuronal processes.

### Gene set enrichment analysis

The Enrichr API in the GSEApy library was utilized to evaluate the functional enrichment of cNMF modules using the top 50 highest-ranking genes from each module, with the complete gene panel serving as the background. We selected terms with an adjusted p-value below 0.05 and a size greater than 2. Additionally, we conducted Gene Set Enrichment Analysis (GSEA) for neuronal and glial subtypes using gene sets from Farsi *et al*., which included the C5 v7.2 collection (comprising 14,765 Gene Ontology terms) from the Molecular Signature Database (http://www.gsea-msigdb.org/gsea/msigdb), SynGO collection, and other literature-derived gene sets. Terms were chosen based on an FDR q-value below 0.05 and meeting expression thresholds.

### Cross-cell-type DEG co-variation analysis

For neuronal, astrocyte, oligodendrocyte, and pericyte DEGs, we computed regional log fold-changes for each gene across 18 molecular tissue regions identified by SPIN. 5 molecular tissue regions (VS, FT, MNG, CTX_HIP, CTX_PIR) were excluded from the analysis due to low neuron abundance. Pearson correlation coefficients were then calculated for every possible cross-cell-type DEG pair, yielding full cross–cell-type correlation matrices for downstream analyses. In parallel, regional neuronal immediate early gene (IEG) score changes were calculated and correlated with glial DEGs to identify glial genes associated with neuronal activity.

### Mapping human GWAS traits with gsMap

gsMap (v1.73.6) was applied to each WT sample to generate single-cell association scores with GWAS traits associated with human schizophrenia, following the GitHub tutorial (https://yanglab.westlake.edu.cn/gps_data/website_docs/html/tutorials.html). Each sample was processed in quick mode using default settings, with an added uniform slice mean to account for multiple biological replicates.

### Purification of synapse fractions

Whole cortices from 4-week and 12-week old male *Grin2a* mutant mice and wild-type littermates (n=5 wild-type, 5 heterozygous knockouts, 5 homozygous knockouts of each age) were used in this study. Mice were euthanized by CO₂ inhalation, after which the cortex was rapidly dissected, flash-frozen in liquid nitrogen, and stored at −80 °C until processing.

Synapse fractions were purified as previously described^68,70^. Briefly, frozen cortex tissue was thawed on ice and dounce-homogenized in ice-cold homogenization buffer (5 mM HEPES pH 7.4, 1 mM MgCl₂, 0.5 mM CaCl₂, supplemented with protease and phosphatase inhibitors). The homogenate was centrifuged at 1,400 g for 10 min at 4 °C, and the resulting supernatant was centrifuged again at 13,800 g for 10 min at 4 °C. The pellet was resuspended in 0.32 M sucrose, 6 mM Tris-HCl (pH 7.5), layered onto a discontinuous sucrose gradient (0.85 M, 1.0 M, and 1.2 M sucrose in 6 mM Tris-HCl pH 7.5), and ultracentrifuged at 82,500 g for 2 h at 4 °C.

The synaptosome fraction, located at the interface between the 1.0 M and 1.2 M sucrose layers, was collected and mixed with an equal volume of ice-cold 1% Triton X-100 (in 6 mM Tris-HCl pH 7.5). After incubation on ice for 15 min, the sample was ultracentrifuged at 32,800 g for 20 min at 4 °C. The resulting pellet was solubilized and resuspended in 1% SDS in H_2_O. This pellet is highly enriched for postsynaptic density proteins, as well as for proteins of the cytomatrix of the active zone (CAZ). Although this fraction has historically been referred to as the “postsynaptic density” fraction, we refer to it as the “synapse fraction” to reflect the abundant presence of CAZ proteins^68^. After solubilization, a small aliquot was removed for protein quantification by BCA assay (Thermo Fisher Scientific), and the remaining material was stored at −80 °C until quantitative mass spectrometry.

### Processing of synapse fractions for mass spectrometry (MS/MS)

The protein in synapse fraction samples from both 4-week and 12-week animals (in 1% SDS) was reduced using 5 mM dithiothreitol and alkylated using 10 mM iodoacetamide at room temperature. The denatured, reduced, alkylated protein samples were then processed using S-Trap sample processing technology (Protifi) following manufacturer’s instructions. The proteins were bound to the S-Trap column via centrifugation and contaminants/detergents were washed away. Sequential digestion steps were then performed on column using 1:20 enzyme to substrate ratio of Lys-C for 2 hours followed by Trypsin overnight at room temperature.

Two Tandem Mass Tag (TMT) 16-plex experiments were constructed, each containing samples from the same age group. Additionally, a pooled reference sample was created using equal amounts of all 30 (15 4-week and 15 12-week) samples to include in each plex for cross-plex comparison. Following digestion, 50 μg of each sample was labeled with TMT16 reagent. Each plex included five wild-type, five *Grin2a*+/-, and five *Grin2a*-/- samples, which were randomly assigned to TMT reporter channels. The last channel 134N contained the pooled reference sample in both plexes. After verifying successful labeling of more than 95% label incorporation, reactions were quenched using 5% hydroxylamine and samples were mixed together. The TMT16 labeled peptides were desalted on a 50 mg tC18 SepPak cartridge and fractionated by high pH reversed-phase chromatography on a 4.6 mm x 250 mm Zorbax 300 extend-c18 column (Agilent). One-minute fractions were collected during the entire elution and fractions were concatenated into 12 fractions for LC-MS/MS analysis.

### Liquid chromatography-Mass Spectrometry analysis (LC-MS/MS)

One microgram of each proteome fraction was analyzed on a QE-HFX mass spectrometer (Thermo Fisher Scientific) coupled to an easynLC 1200 LC system (Thermo Fisher Scientific). Samples were separated using 0.1% Formic acid / 3% Acetonitrile as buffer A and 0.1% Formic acid / 90% Acetonitrile as buffer B on a 27cm 75um ID picofrit column packed in-house with Reprosil C18-AQ 1.9 mm beads (Dr Maisch GmbH) with a 110 min gradient consisting of 2-6% B in 1 min, 6-20% B in 62 min, 20-30% B for 22 min, 30-60% B in 9 min, 60-90% B for 1 min followed by a hold at 90% B for 5 min. The MS method consisted of a full MS scan at 60,000 resolution and an AGC target of 3e6 from 350-1800 m/z followed by MS2 scans collected at 45,000 resolution with an AGC target of 1e5 with a maximum injection time of 105 ms and a dynamic exclusion of 15 seconds. The isolation window used for MS2 acquisition was 0.7 m/z and 20 most abundant precursor ions were fragmented with a normalized collision energy (NCE) of 29 optimized for TMT16 data collection.

### Database search and MS/MS quantification

The data was analyzed using Spectrum Mill MS Proteomics Software (Broad Institute) with a mouse database from Uniprot.org downloaded on 04/07/2021 and containing 55,734 entries. Proteins identified by one or more peptides were selected for further analysis. Search parameters included: ESI Q Exactive HCD v4 35 scoring parent and fragment mass tolerance of 20 ppm, 40% minimum matched peak intensity, trypsin allow P enzyme specificity with up to four missed cleavages, and calculate reversed database scores enabled. Fixed modifications were carbamidomethylation at cysteine. TMT labeling was required at lysine, but peptide N termini could be labeled or unlabeled. Allowed variable modifications were protein N-terminal acetylation, deamidation and methionine oxidation. Protein quantification was achieved by taking the ratio of TMT reporter ions for each sample over the TMT reporter ion for the pooled reference channel. TMT16 reporter ion intensities were corrected for isotopic impurities in the Spectrum Mill protein/peptide summary module using the afRICA correction method which implements determinant calculations according to Cramer’s Rule and correction factors obtained from the reagent manufacturer’s certificate of analysis (https://www.thermofisher.com/order/catalog/product/90406) for lot numbers VH310017. After performing median-MAD normalization, a moderated two-sample t-test was applied to the datasets to compare wild-type, *Grin2a*+/-, and *Grin2a*-/- sample groups at each age. A comprehensive list of differentially abundant proteins for 3-month *Grin2a*+/- and *Grin2a*-/- cortical synapse fraction samples is provided in Supplementary Table S6.

## Data and code availability

The datasets and codes used in our study will be made public upon publication of the manuscript.

## Acknowledgements

We thank S. McCarroll for providing valuable insights into data interpretation, Z. Tang and K. Maher for assistance on computational analysis, and J. Tian, S. Furniss, and C. Mehaffey for suggestions on manuscript writing. X.W. acknowledges the support from Stanley Center for Psychiatric Research at Broad Institute, Packard Fellowship for Science and Engineering, Merkin Institute Fellowship, National Institutes of Health (NIH) DP2 New Innovator Award (1DP2GM146245) and NIH BRAIN CONNECTS (UM1 NS132173). W.X.W. is a Damon Runyon–National Mah Jongg League Fellow, supported by the Damon Runyon Cancer Research Foundation (DRG no. 2512-23).

## Author information

These authors contributed equally: Mingrui Wu, Jiahao Huang

Contributions: M.W. designed and performed STARmap and RIBOmap experiments. J.H. performed downstream imaging data processing. J.H. and M.W. performed computational analysis. S.A., K.B., M.Y., B.D., H.K., and S.A.C. performed synaptic proteome profiling. I.P. and Y.Z. prepared mouse brain samples. H.C. prepared STARmap and RIBOmap probes. Z.F., S.L., and W.X.W. offered inputs on data analysis and interpretation. All authors contributed to writing and revising the manuscript and approved the final version. X.W. and M.S. conceptualized and supervised the project.

Corresponding authors: Xiao Wang, Morgan Sheng

## Competing interests

X.W. is a scientific cofounder and consultant of Stellaromics and Convergence Bio. M.S. is SAB member and/or consultant of Neumora, Biogen, Illimis, CurieBio, Astex. Other authors declare no competing interests.

---

**Extended Data Fig. 1:**
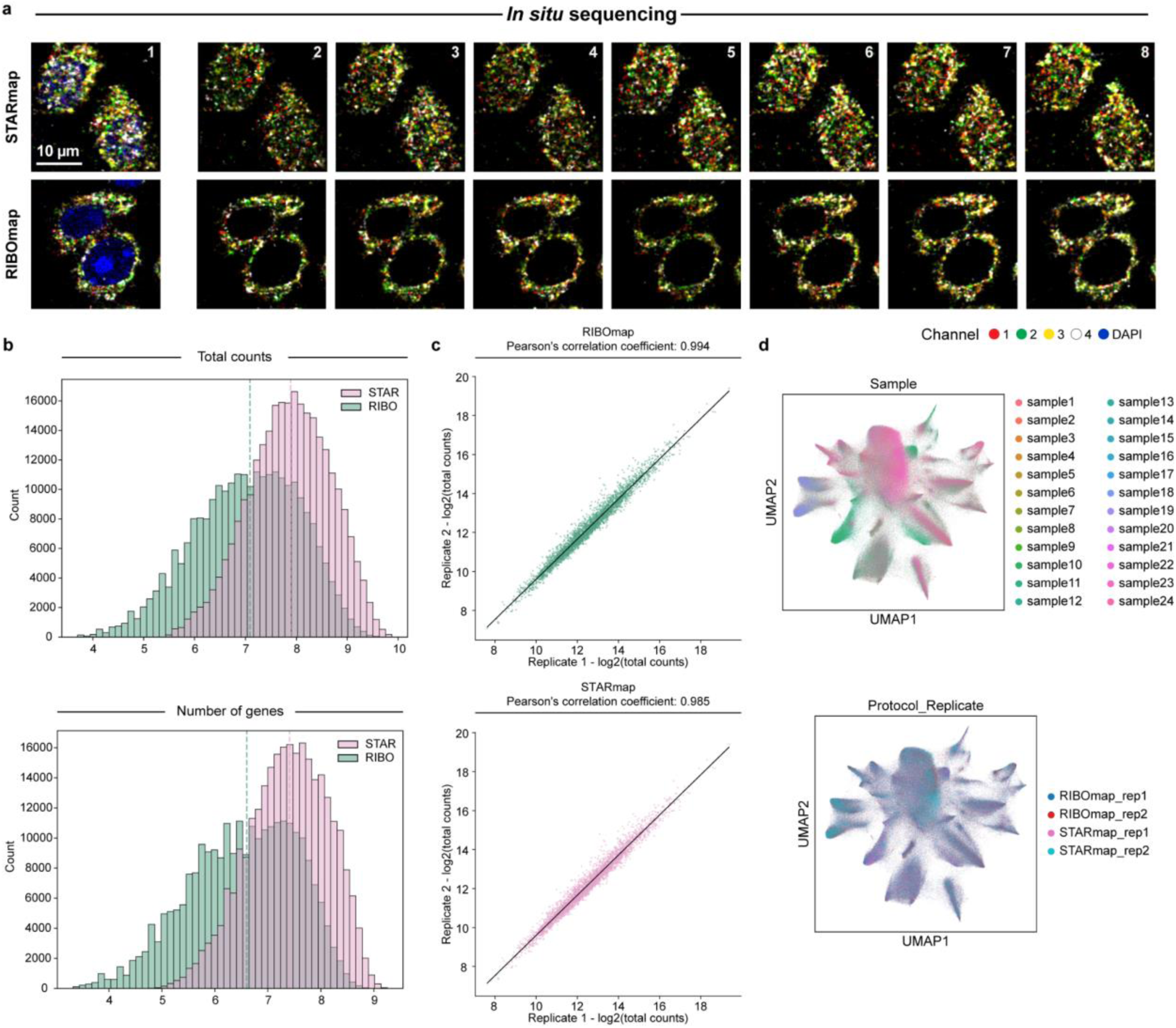
Quality assessment of spatially resolved single-cell transcriptional and translational data. **a**, Representative images showing the sequential mapping of cDNA amplicons for STARmap (top) and RIBOmap (bottom) in mouse brain tissue sections. **b**, Histogram of the number of transcripts (top) and genes (bottom) in each cell after logarithmic transformation (log2) of STARmap and RIBOmap datasets. The median reads per cell are 237 for STARmap and 136 for RIBOmap, while the median genes per cell are 170 for STARmap and 97 for RIBOmap. **c**, The correlation of the number of reads for each gene between the two biological replicates in RIBOmap (top) and STARmap (bottom). **d**, UMAP showing the distribution of cells across 24 samples (top) and replicates in each modality (bottom).

**Extended Data Fig. 2:**
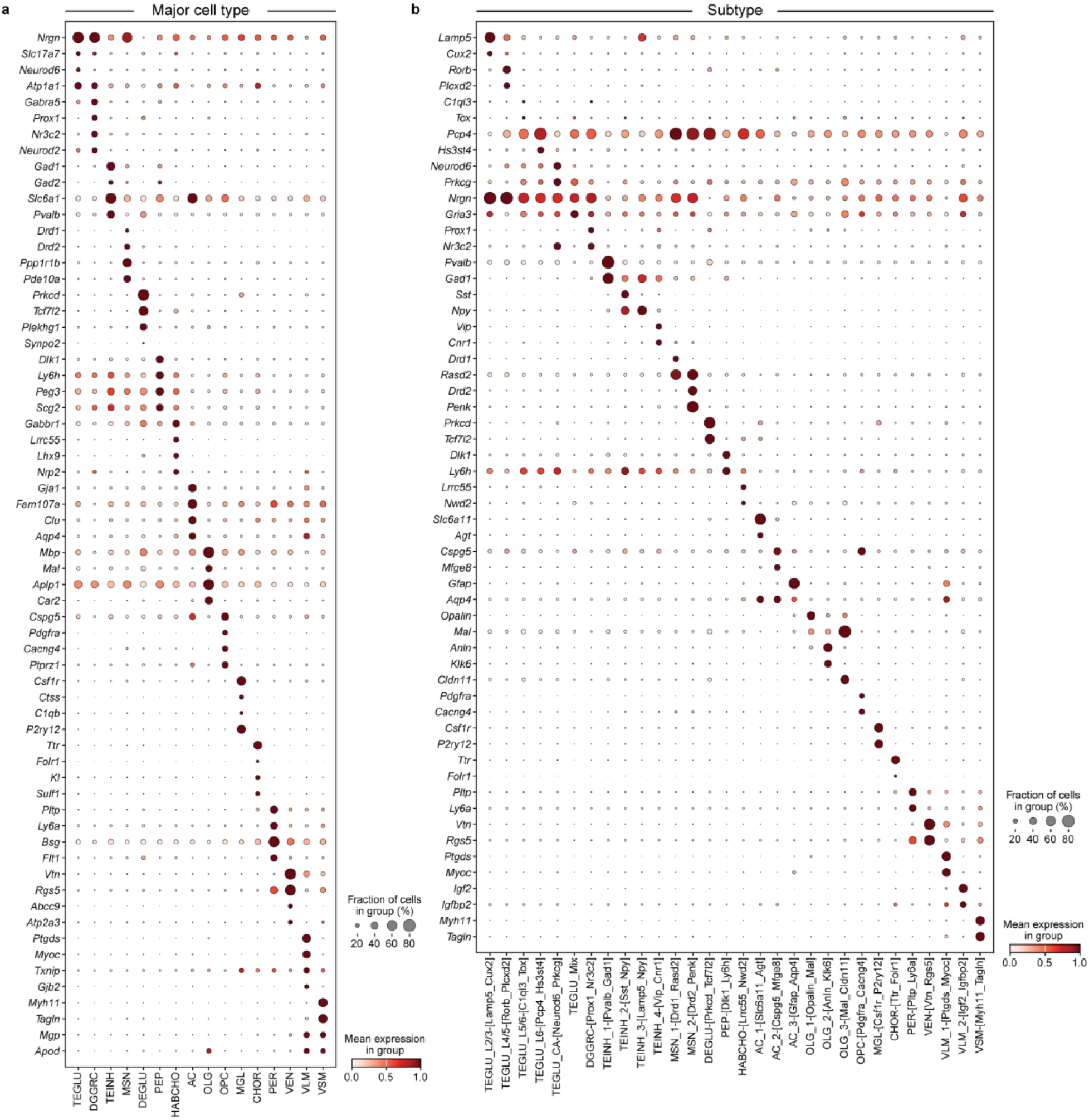
Marker genes for major cell types and subtypes on the integrated dataset. **a**, Dot plot showing the expression level of representative markers across different major cell types. Color scale, averaged gene expression; dot size, the percentage of cells expressing the genes within each major cell type. **b**, Dot plot showing the expression level of representative markers across different subtypes. Color scale, averaged gene expression; dot size, the percentage of cells expressing the genes within each subtype.

**Extended Data Fig. 3:**
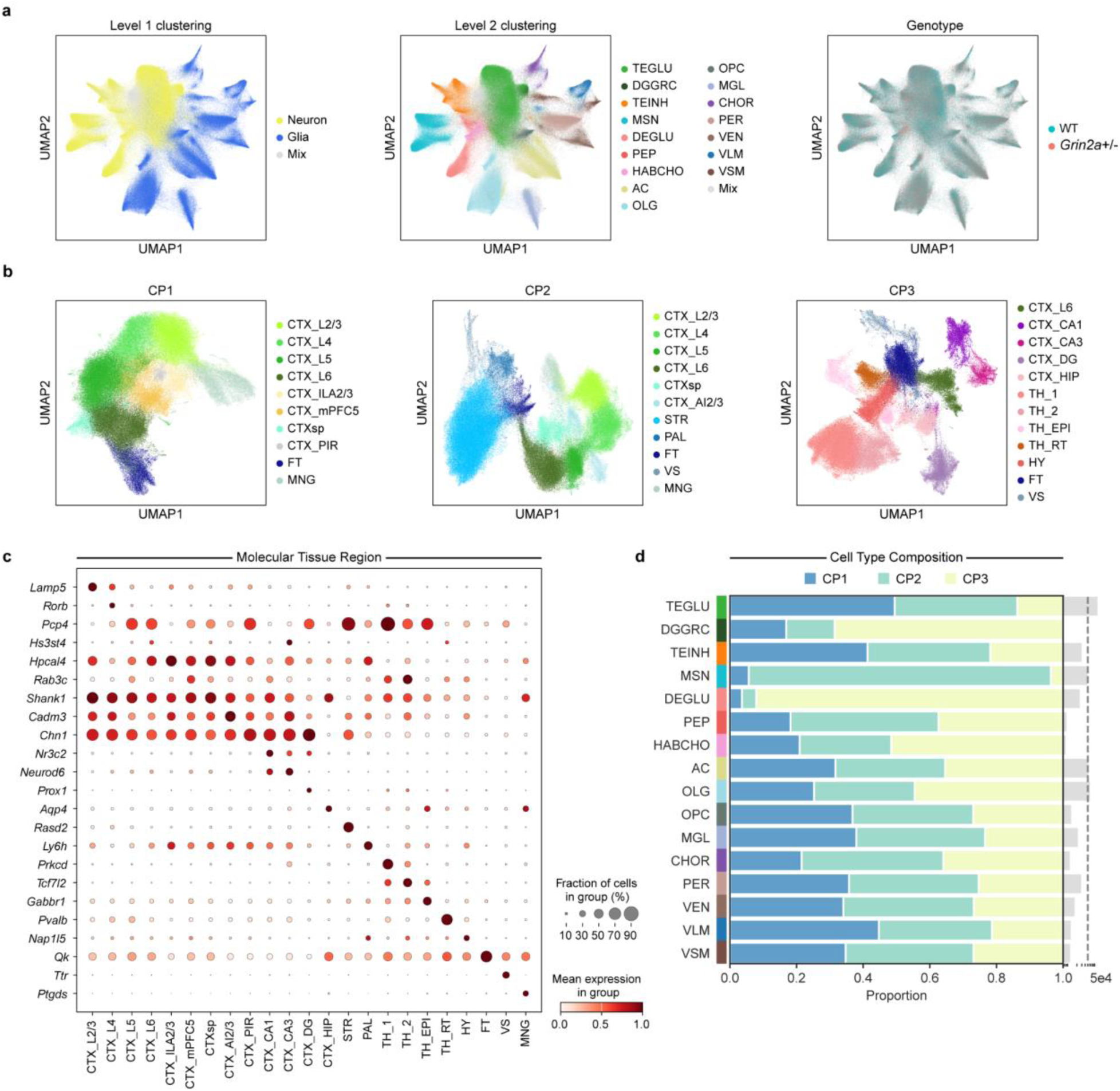
Multi-level cell type classification and tissue region identification with SPIN. **a**, UMAPs showing the distribution of cells colored by level 1 cell type annotations (left), level 2 cell type annotations (mid), and genotypes (right). **b**, UMAP of molecular tissue region annotation by SPIN at each coronal position. **c**, Dot plot showing the expression level of representative markers across molecular tissue regions. Color scale, averaged gene expression; dot size, the percentage of cells expressing the genes within each region. **d**, Bar plot showing major cell type composition at each coronal position.

**Extended Data Fig. 4:**
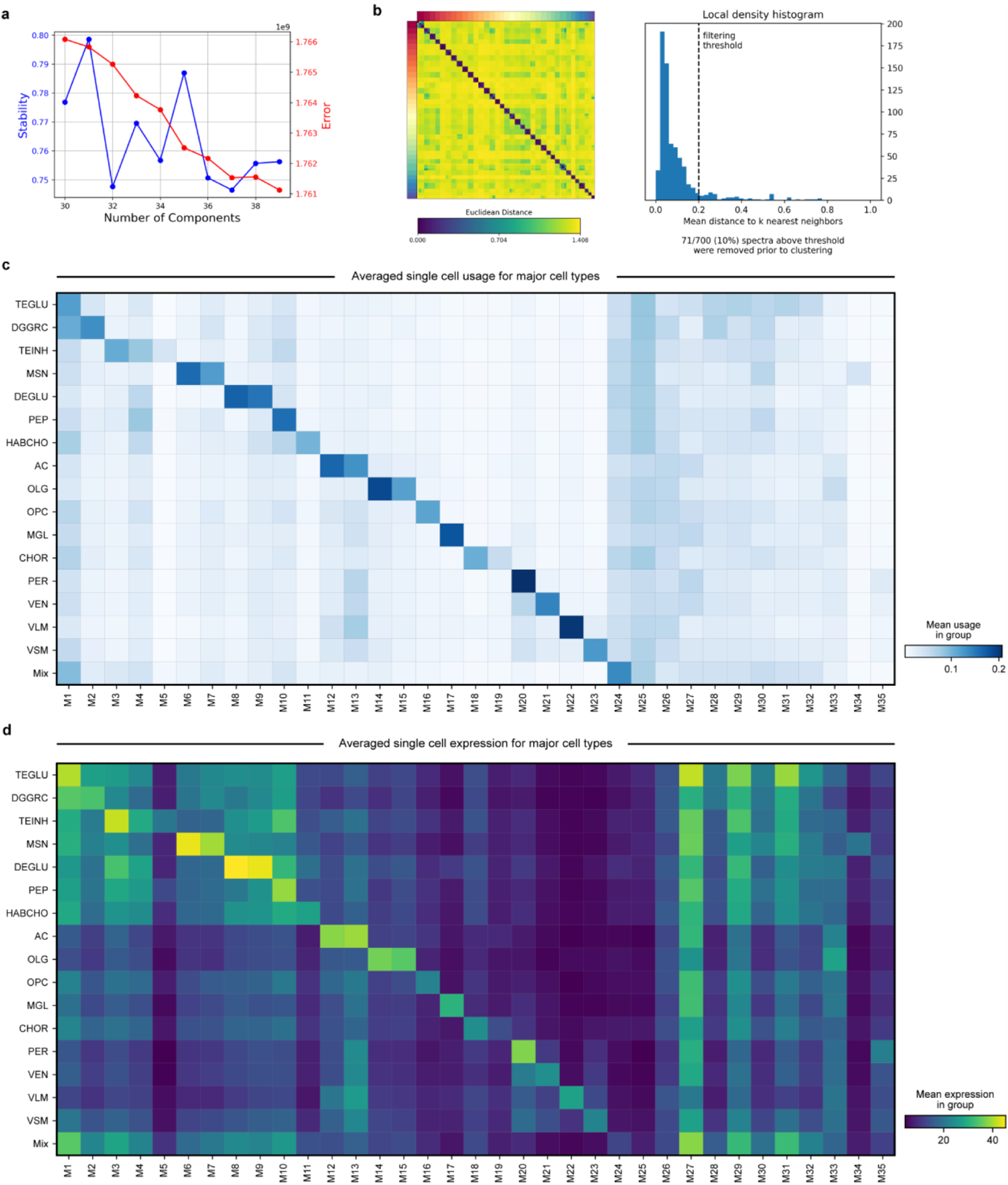
Gene module identification with cNMF. **a**, Visualization showing the tradeoff between error and stability of cNMF factors in relation to the number of factors k. 35 factors were requested based on these results. **b**, Clustermap showing the consensus matrix factorization estimates, with each color on the x- and y-axes representing one of the 35 cNMF factors. **c-d**, Matrix plot showing single cell usage score (**c**) and averaged gene expression (**d**) of each gene module across major cell types.

**Extended Data Fig. 5:**
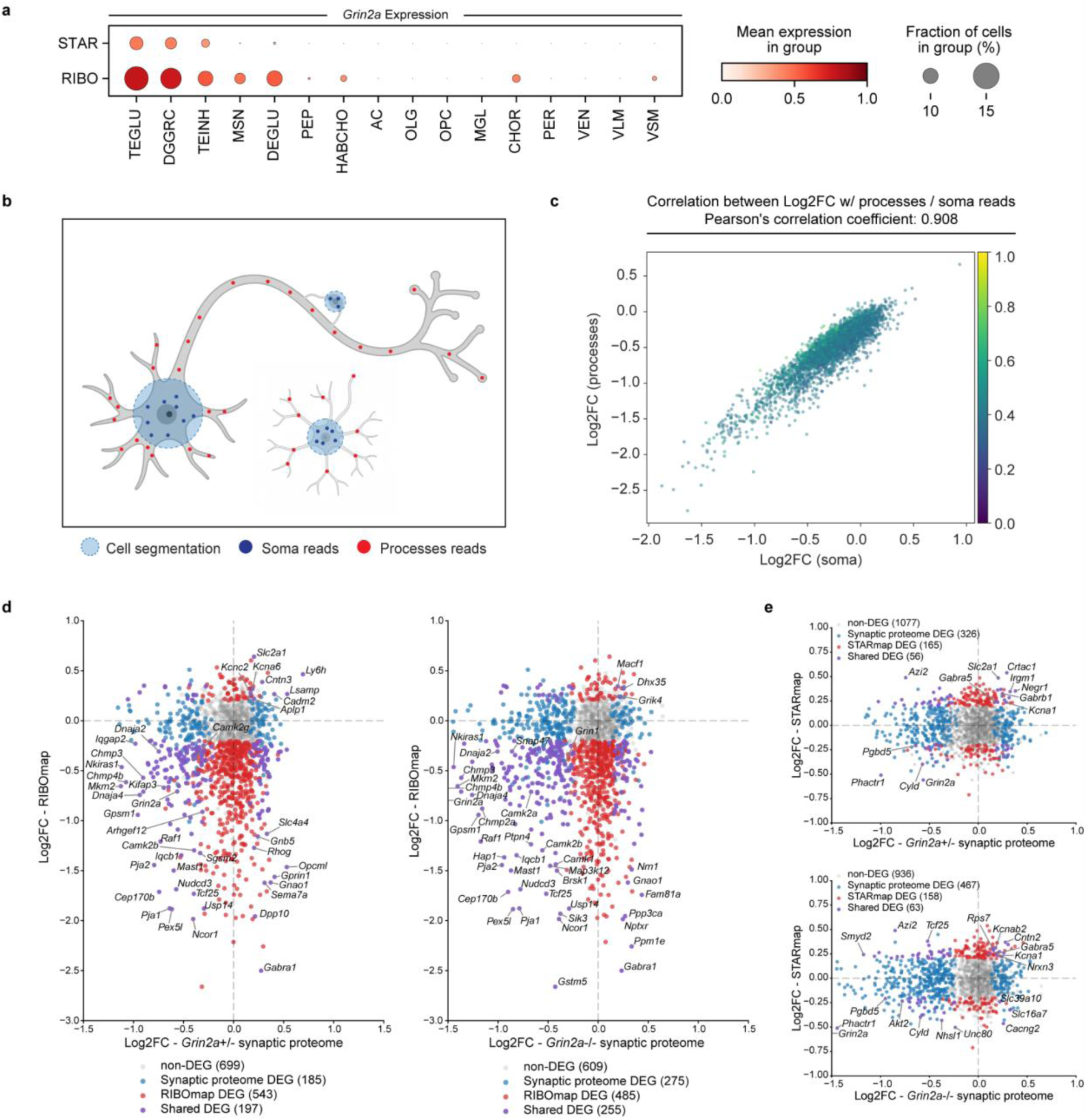
Comparison between RIBOmap/STARmap and synaptic proteome. **a**, Dot plot showing the expression of *Grin2a* in major cell types measured by STARmap and RIBOmap. Color scale, averaged gene expression; Dot size, expression percentage. **b**, Schematics showing soma-vs. processes-reads at the single cell level. Signals outside the segmentation area were treated as reads in the neuronal processes. **c**, Scatter plot comparing the log fold-change values of each gene calculated from soma-reads versus process-reads. **d**, RIBOmap log fold-change vs. cortical synaptic proteome log fold-change in *Grin2a*+/- and *Grin2a*-/- mutants. **e**, STARmap log fold-change vs. cortical synaptic proteome log fold-change in *Grin2a*+/- and *Grin2a*-/- mutants.

**Extended Data Fig. 6:**
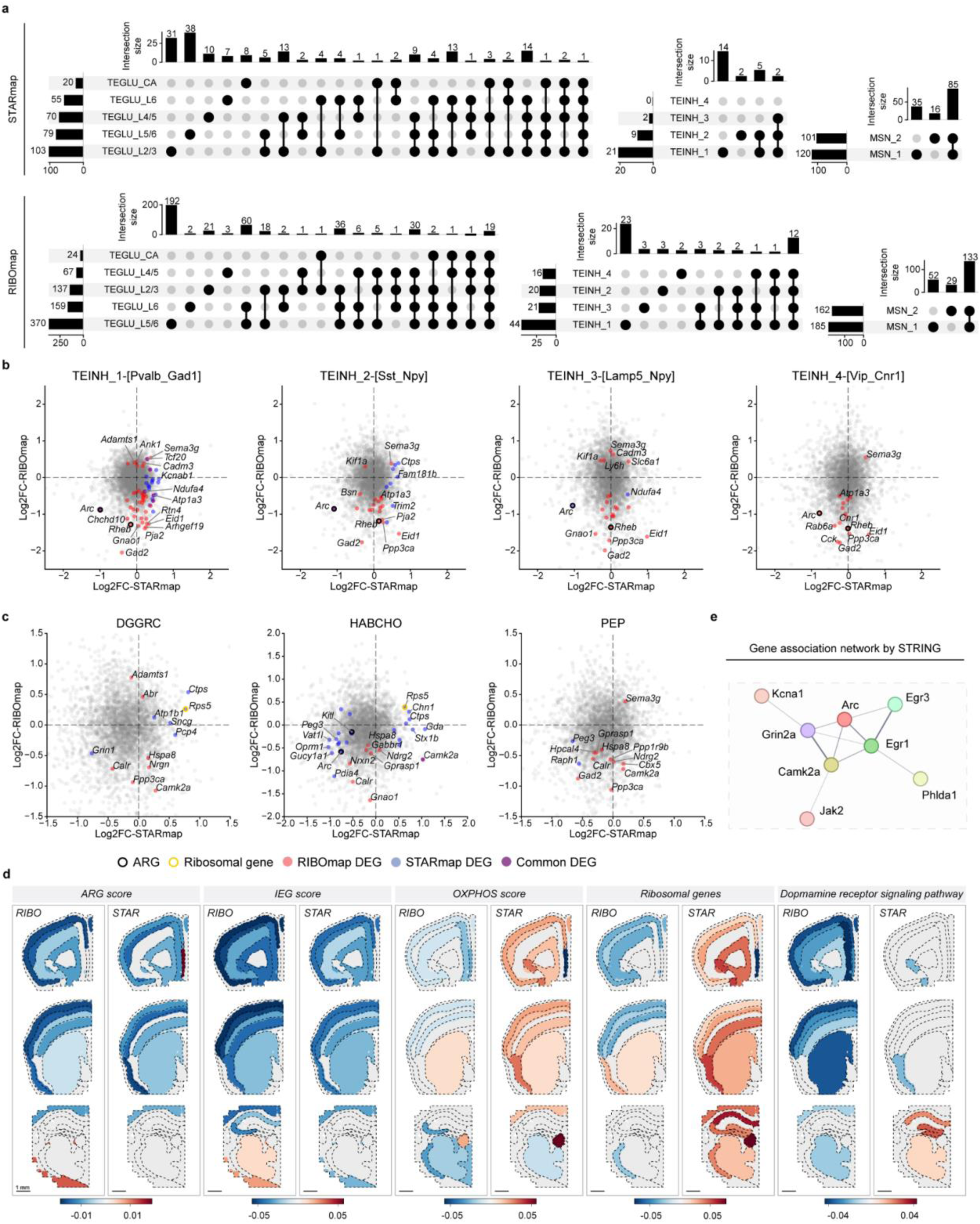
Neuronal subtype DEG. **a**, STARmap and RIBOmap DEG overlap between TEGLU, TEINH, and MSN subtypes. **b**, STARmap vs. RIBOmap log fold-change plot of 4 TEINH subtypes. **c**, STARmap vs. RIBOmap log fold-change plot of DGGRC, HABCHO, and PEP. **d**, Spatial visualizations of gene set score differences between two genotypes in STARmap and RIBOmap. **e**, Pathway-level interactions among genes showing *Grin2a*-dependent translation reduction by the STRING interactome database. Edge width represents association confidence.

**Extended Data Fig. 7:**
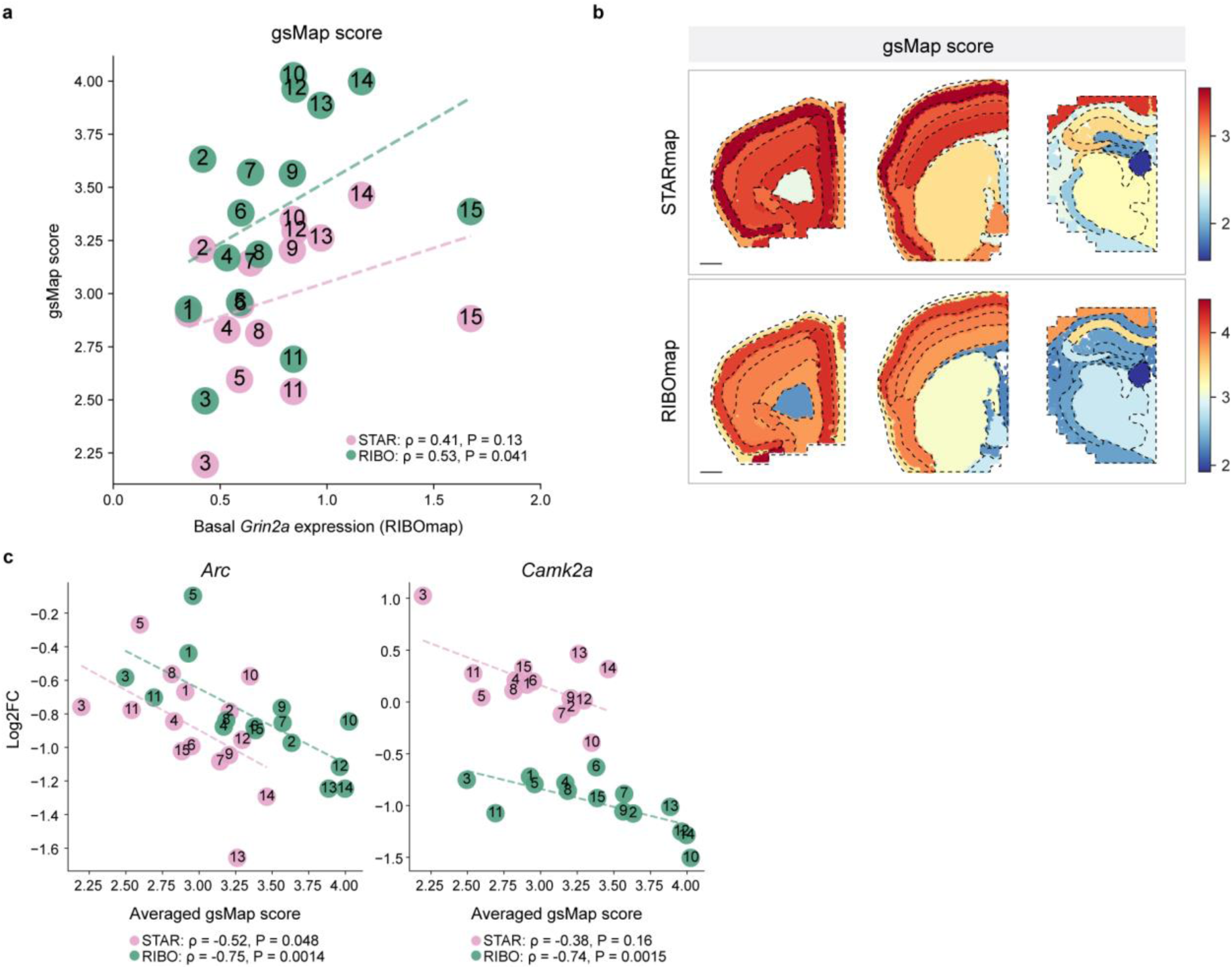
Mapping human GWAS traits to STARmap/RIBOmap data with gsMap. **a**, Scatter plot showing the correlation between basal RIBOmap *Grin2a* expression level and gsMap score prediction across neuronal subtypes. **b**, Spatial heterogeneity of averaged gsMap score in STARmap (top) and RIBOmap (bottom) across tissue regions. Scale bar, 1mm. **c**, Relationship between the gsMap score and the RIBOmap log fold-change of example genes across neuronal subtypes.

**Extended Data Fig. 8:**
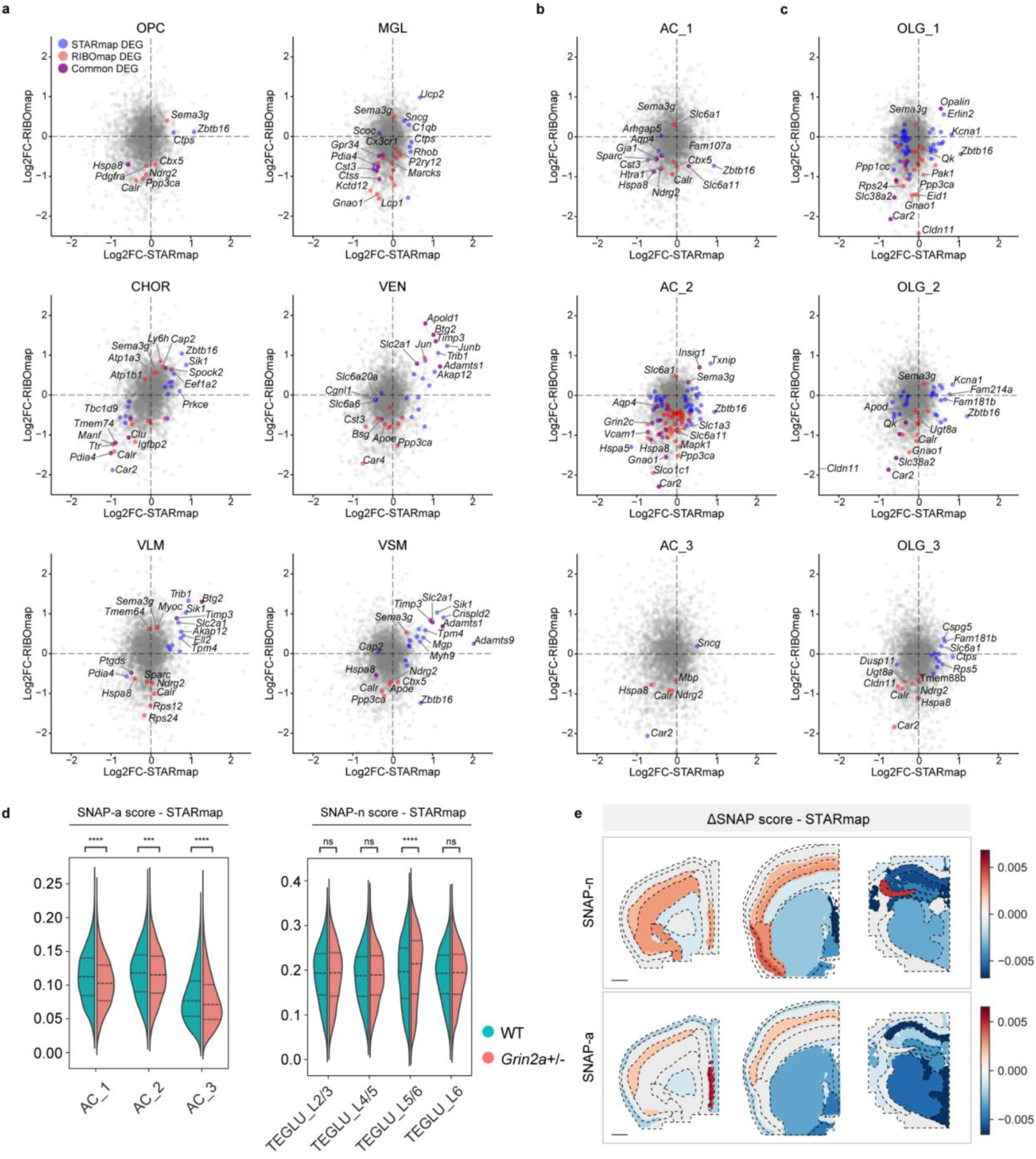
Non-neuronal cell type DEG. **a**, STARmap vs. RIBOmap log fold-change plot of 6 remaining non-neuron major cell types. **b-c**, STARmap vs. RIBOmap log fold-change plot of astrocyte (**b**) and oligodendrocyte (**c**) subtypes. **d**, Violin plots illustrating the difference between WT and *Grin2a*+/- in SNAP-a gene program scores in astrocyte subtypes (left) and in SNAP-n gene program scores in cortical TEGLU subtypes (right) in STARmap. P values were calculated using a two-sided independent t-test. ****P < 0.0001. **e**, Spatial visualizations of SNAP-n score difference in cortical neurons (top) and SNAP-a score difference in astrocytes (bottom) in STARmap. Scale bar, 1mm.

**Extended Data Fig. 9:**
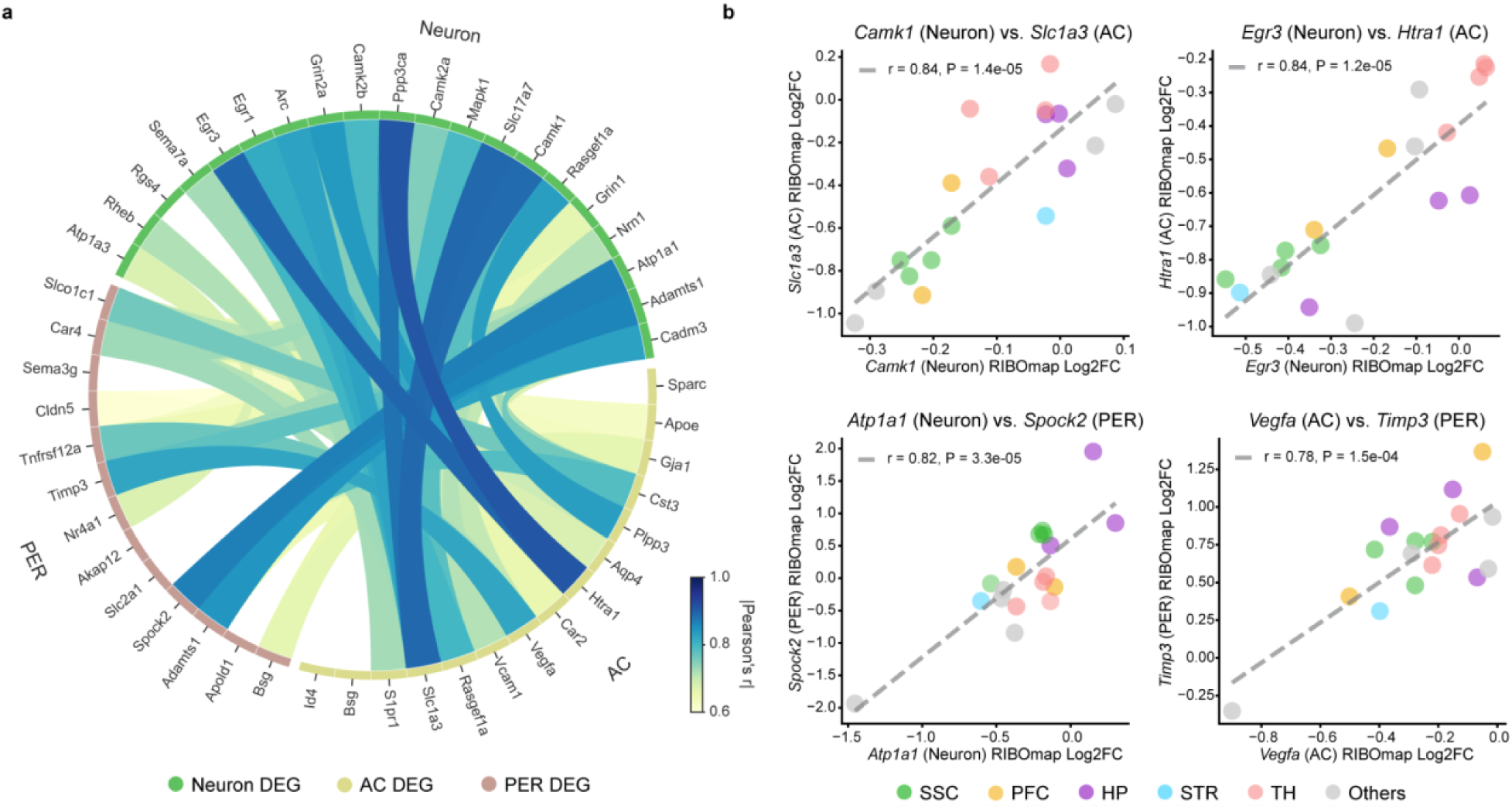
NVU-associated cross-cell-type DEG co-variation. **a**, Chord diagram showing pairwise cross-cell-type NVU-associated DEG correlations of region-specific RIBOmap log fold-change in neuron, astrocyte, and pericyte. Chord color, absolute value of pairwise Pearson’s correlation coefficient r. **b**, Relationship between RIBOmap log fold-changes of example cross-cell-type DEG pairs.

## Notes

### Summary of Updates

Author information updated: co-first author and corresponding author labels

